# ChromDL: A Next-Generation Regulatory DNA Classifier

**DOI:** 10.1101/2023.01.27.525971

**Authors:** Christopher Hill, Sanjarbek Hudaiberdiev, Ivan Ovcharenko

**Affiliations:** Computational Biology Branch, National Center for Biotechnology Information, National Library of Medicine, National Institutes of Health, Bethesda, MD, 20892, USA; School of Engineering and Applied Science, University of Pennsylvania, Philadelphia, PA, 19104, USA

## Abstract

**Motivation:** Predicting the regulatory function of non-coding DNA using only the DNA sequence continues to be a major challenge in genomics. With the advent of improved optimization algorithms, faster GPU speeds, and more intricate machine learning libraries, hybrid convolutional and recurrent neural network architectures can be constructed and applied to extract crucial information from non-coding DNA.

**Results:** Using a comparative analysis of the performance of thousands of Deep Learning (DL) architectures, we developed ChromDL, a neural network architecture combining bidirectional gated recurrent units (BiGRU), convolutional neural networks (CNNs), and bidirectional long short-term memory units (BiLSTM), which significantly improves upon a range of prediction metrics compared to its predecessors in transcription factor binding site (TFBS), histone modification (HM), and DNase-I hypersensitive site (DHS) detection. Combined with a secondary model, it can be utilized for accurate classification of gene regulatory elements. The model can also detect weak transcription factor (TF) binding with higher accuracy as compared to previously developed methods and has the potential to accurately delineate TF binding motif specificities.

**Availability:** The ChromDL source code can be found at https://github.com/chrishil1/ChromDL.

## Introduction

The availability of large genome sequencing datasets has led to the development of numerous computational algorithms for the prediction of chromatin features and gene regulatory elements (referred to as regulatory features here) in the human genome in the past two decades (1; 2; 3; 4; 5; 6; 7) In particular, the advent of DL algorithms has been shown to be effective in the prediction of DHSs, HMs and TFBSs (8; 9; 10; 11) The neural network architectures commonly take genomic sequences as input and apply a series of non-linear transformations before computing probability scores for a series of target labels, each corresponding to a specific regulatory feature. These prediction efforts have become increasingly feasible to create algorithms for with the improvement of machine learning libraries, increased GPU speeds, and sophisticated optimization algorithms, all of which yield convergence to a high-performing solution in a fraction of time compared to earlier generations of neural networks.

One of the first-generation regulatory feature prediction models was DeepSEA (12), which was constructed using three CNNs (Fig S1). DeepSEA paved the way for next generation neural networks in attempting to improve prediction accuracy of TF binding, open chromatin, and HMs (919 regulatory features in total) and allowed the prioritization of expression quantitative trait loci (eQTLs) and disease associated variants (12).

Subsequent models investigated the effectiveness of recurrent neural networks (RNNs) in addition to CNNs (8; 9; 10; 11). These RNN layers share parameters between their units as CNNs do, but also contain internal memory components that allow them to “remember” previous passes over the input. RNNs are generally considered to be more powerful when dealing with sequential data processing, whereas CNNs are typically used to capture recurring patterns and motifs within the whole input data. Since the sequences fed into these models are represented as one-hot encoded matrices, two RNN classes can be combined in a larger architecture to improve performance when processing the inputs: Long Short-Term Memory (LSTM) and Gated Recurrent Units (GRUs) that act in a similar fashion and often improve performance as compared to traditional RNNs (9; 13; 14; 15).

While both RNNs and LSTMs contain feedback loops that help them “remember” the previous iteration, LSTMs contain an additional memory unit, which gives them more control about when information is remembered, when it is utilized, and when it is forgotten, making them more flexible than a traditional RNN (14; 15). GRUs conversely contain an update and a reset gate that control what information is retained but lack an output gate, instead feeding the input and the reset gate back into the update gate. Both layers attempt to combat the vanishing gradient problem, a well-documented issue that prevents weights from changing when gradients become increasingly smaller with each epoch of training in traditional RNNs (16).

Another feature of these RNN classes is their ability to be bidirectional, where two identical hidden layers read the input in a forward and reverse direction simultaneously to give the model context of the past, present, and future input (14; 15; 17). Two models introduced by Quang *et al*. (18), denoted as DanQ and DanQ-JASPAR, found success with these bidirectional RNNs as well as initializing their model with motifs from the JASPAR database (19). The DanQ neural network was constructed with one convolutional layer that mirrored DeepSEA’s input processing, followed by a BiLSTM layer, before using DeepSEA’s exact dense and output layer architecture, subsequently improving performance metrics (Fig S2). DanQ-JASPAR increased the number of filters in the convolutional layer and number of units in the BiLSTM, and together with the initialized motifs observed improved performance. In this way, the authors proved that BiLSTMs could be used in regulatory feature prediction contexts using only DNA sequences as input, and opened up the possibility of combining CNNs and RNNs in further intricate architectures to improve performance metrics.

Here, we propose ChromDL, a neural network architecture that improves on previously developed models using a combination of CNNs and RNNs. ChromDL combines BiGRU layers, BiLSTM layers, and CNN layers in an architecture that differs quite substantially from both DeepSEA and DanQ. ChromDL surpasses these methods in accuracy of regulatory feature prediction. The model also detects a significantly higher proportion of weak TFBS ChIP-seq peaks and demonstrates the potential to more accurately predict TF binding affinities. We also show that the TREDNet DL second-phase enhancer classifier (20) built on top of ChromDL is capable of detecting regulatory variants according to validation using reporter assay quantitative trait loci (raQTLs). We believe ChromDL will allow accurate predictions of gene regulatory mechanisms, modeling gene regulatory networks, and detection of disease-causative non-coding variants with utmost precision.

## Methods

### Model Architecture and Training

ChromDL is a DL neural network containing two internal CNNs and three bidirectional RNNs. The neural network in its entirety extracts sequence features using weight matrices in sequential layers to minimize errors in prediction of the outputs, in this case being the 919 regulatory feature labels.

This model contains eleven layers, with ten of these being distinct (Fig 1). The 1kbp input sequence, encoded in one-hot representation (1000 × 4), was fed into a 128 unit BiGRU layer, which was then fed into a separable convolutional layer with 750 filters and a kernel size of 16, followed by a standard convolutional layer with 360 filters and a kernel size of 8. Both CNN layers had L1/L2 regularization of 1E-08 and 5E-07, respectively, and rectified linear unit (ReLU) activation before being maximally pooled with a 1×4 window. Next, the data was fed into a 128 unit BiLSTM layer with 20% dropout and applied batch normalization, before an average pooling layer with a 1×8 window. Finally, a second 128 unit BiLSTM layer was applied before the data was flattened and fed into the 919 length dense layer with sigmoid activation that produced the scores for all of the labels included in the dataset. In total, the model contained 10,414,957 parameters, with 512 non-trainable parameters. The exact technical specifications of these layers can be found in the Supplemental Materials (see *ChromDL Layer Specifications*).

**Fig. 1:**
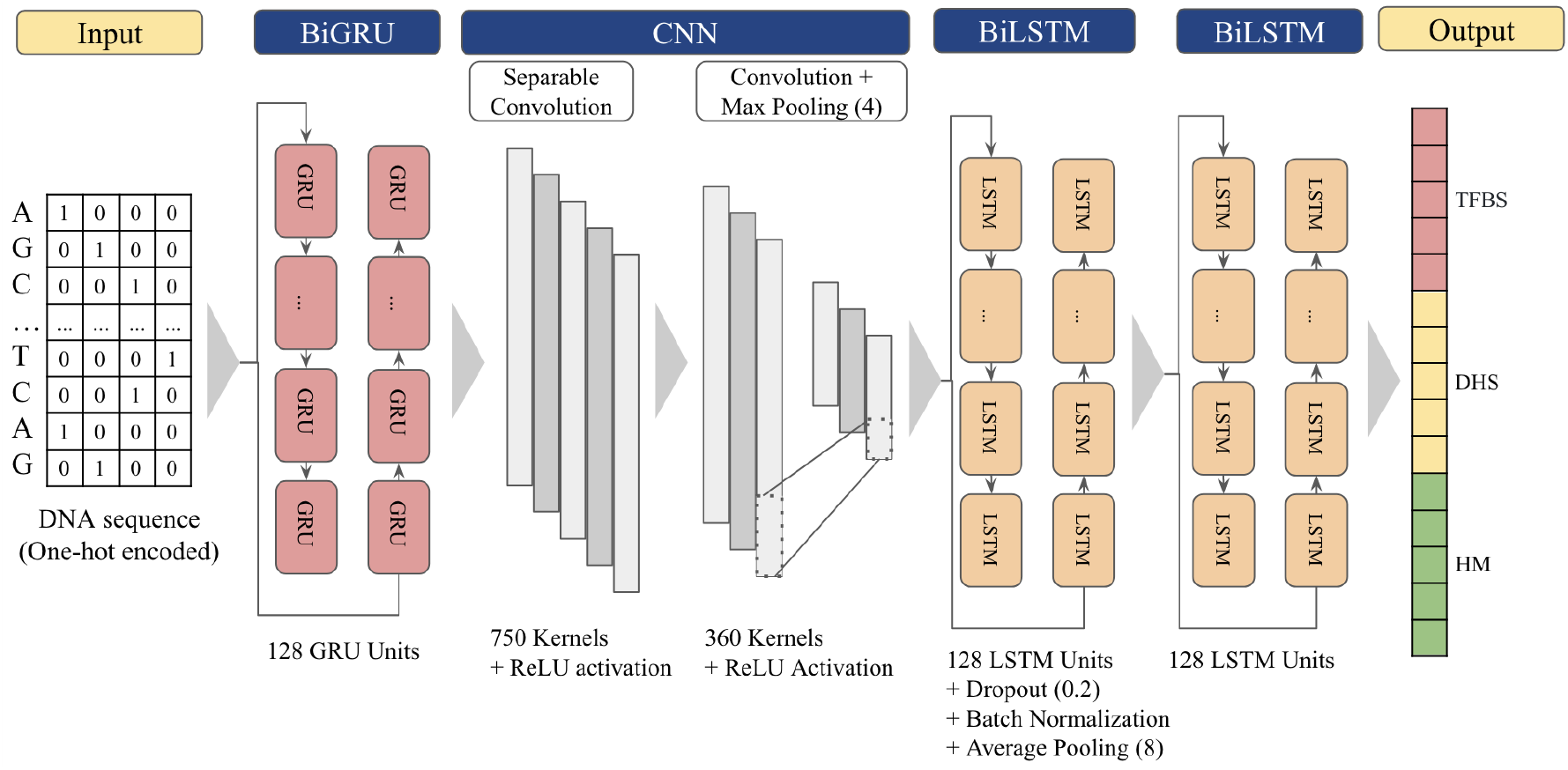
Schematic representation of ChromDL’s deep learning architecture.

This model utilized the same dataset and target feature labels described in DeepSEA in it’s training, testing, and validation (12) (Table S9). Briefly, the human GRCh37 reference genome was split into 200-bp bins, and each TF-bound bin was labeled with 919 regulatory features. These labels include 690 TFBS, 125 DHS, and 104 HM regions. Each 200-bp bin included in the dataset was extended to 1kbp to include the flanking regions before being fed to the models in one-hot representation. 4,000 regions from chromosome 7 were used as the validation set, the entirety of chromosomes 8 and 9 as the test set, and the rest of the autosomal chromosomes as the training set. All available training/validation regions were included in training this model, and all available testing regions were utilized for analyzing performance. The models were evaluated using auROC and auPRC metrics. The average of the model outputs for forward and reverse strand sequences of the input regions were used as probabilities and compared to the true labels.

To train this model, we minimized the binary cross-entropy loss function between the true labels and the predicted labels after the sequences were processed by the model. This loss function can be expressed as:

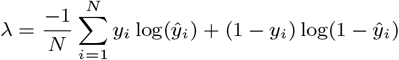

where N is the output size (the 919 predicted labels), *ŷ*_*i*_ is the ith scalar value in the model’s output, and *y*_*i*_ is the corresponding target value.

The Adaptive Moment Estimation (Adam) optimizer was ultimately chosen in the training of the model due to it’s properties of both the Adagrad and RMSprop optimizers, and it’s relatively rapid convergence to solutions (21; 22) (Table S1). The optimizer utilizes a stochastic gradient descent method that is based on the adaptive estimation of first and second order moments. We trained our models using the Adam optimizer with a learning rate of 1E-03 and the parameters *β*_1_ = 0.9, *β*_2_ = 0.999, *ϵ* = 1E-08 (23; 24). A minibatch of size 500 was utilized in each training step.

The models were trained for 100 epochs or until stagnation of validation loss values. ChromDL models stagnated within the first 20 epochs. Each epoch took approximately 16 hours on a single core NVIDIA Tesla K80 GPU processor with 24 GB RAM. All our models were implemented in Tensorflow v2.4.0 with built-in Keras libraries (24).

### TF ChIP-seq signal intensity analysis

For this analysis, we collected signal values from the ChIP-seq peak files of the TFBS 690 datasets to compare correlations between these values and ChIP-seq predictions for four models: ChromDL, DeepSEA, DanQ, and DanQ-JASPAR. We limited our investigation to chromosomes 8 and 9, as the model had been trained and validated using the TFBS ChIP-seq peak data from all other autosomes. Ten equal percentile bins were created from the signal values of the individual peaks for each cell line. For every peak in the dataset, the center of the peak was taken and flanked with 500bp windows on either side to obtain a 1kbp sequence which was then fed through the respective model. The prediction value for each peak was isolated from the 919-output label vector and paired with the signal values from the published data. The 3%, 5%, and 10% false positive rate (FPR) thresholds were calculated for each model using the original DeepSEA testing dataset and used to determine whether a given peak was correctly predicted for the corresponding TFBS label. These proportions predicted correctly were pooled for all 690 cell lines for each of the ten bins and plotted for each of the four models. To measure the significance of the pooled data, three separate one sided independent t-tests were conducted in each percentile bin using the SciPy stats library to calculate the p-value of ChromDL having a higher average proportion than each of the three other models of interest (25). P-values *<* 1E-05 were denoted as significant.

### Motif/Co-Factor performance

We utilized two separate tools for the discovery of motifs in the transcription factor binding site ChIP-seq peak files: the HOMER Motif Analysis tool and the MEME-ChIP Motif Analysis of Large Nucleotide Datasets (26; 27; 28). Before these tools were used, the peak regions that were predicted correctly in the previous analysis were extracted for each of the four models (ChromDL, DeepSEA, DanQ, DanQ-JASPAR). The two motif tools were then run on the set of predicted positive peaks using a 3% FPR in motifs where Protein Binding Microarray (PBM) data was available as found in their ChIP-seq files (not extended to 1kbp). For MEME-ChIP, the peaks were converted to fasta format, and the command line interface was used to detect the top 20 motifs, as any motifs outside the top 20 were generally found to be noisy and insignificant. The script utilized an external control file as the background that was assembled by merging all the TF ChIP-seq narrow peak regions from ENCODE of HepG2, K562, and H1 cell-lines, with an assumption that these are empirically observed TF-binding regions (1; 2). For each trial, any overlaps in the control file with the target regions were removed from the background.

The HOMER and MEME-ChIP *de novo* generated motifs were compared to published PBM motif logos and position weight matrices (PWMs) through each tool’s suggested motif matches and visual analysis. The TOMTOM Motif Comparison Tool was used to generate p-values in attempting to map the *de novo* motif PWMs to the PBM published PWMs (29).

This motif analysis was run for the following cell lines with available PBM data: BCL11A, c-Fos, c-Jun, and TBP (30; 31; 32). The resulting motif logos for the two BCL11A cell lines and five TBP cell lines in the DeepSEA dataset were inconclusive and could not be mapped to published PBM PWMs. All motif PWMs and logos were pulled from the Harvard Universal PBM Resource for Oligonucleotide Binding Evaluation (UniProbe) Database, which consolidated the data from various studies that looked at these cell lines individually (33).

### Enhancer Scoring Validation

For our enhancer validation, we utilized the CNN architecture from the TREDNet DL second-phase enhancer classifier and attached it to our regulatory feature model (20). This classifier took the 919 output labels from our feature prediction model and produced a single output enhancer prediction score. This model’s architecture contained three CNN layers of increasing filter size with ReLU activation, a 180-unit dense layer, a batch normalization layer, two max pooling and dropout layers, and a LeakyReLU activation function before the single neuron dense output layer (Fig S3).

The mutagenesis for the enhancer validation was conducted using the binding sites of EP300 (optimal IDR peaks of the ENCODE ChIP-seq were used, 5,526 peaks in total) and inputting 4,000 regions for each individual enhancer. Each region was in silico mutated by looking at one bp of the 1kbp in the sequence and generating the four alternate alleles that could be recognized by the model (A, C, G, T, and N), with the rest of the sequence remaining unchanged. The one-hot encoded representations of these 4,000 sequences were run through ChromDL and this secondary enhancer model, and the scores were then compared with the wildtype enhancer score.

We then established eight enhancer definitions for the training of this model: 1) ChromHMM Strong Enhancer States, 2) ChromHMM Weak Enhancer States, 3) H3K27ac peaks, 4) H3K4me1 peaks, 5) Overlapping regions between H3K27ac and H3K4me1 peaks, 6) Overlapping regions between H3K27ac and H3K4me1 with any H3K4me3 peaks removed, 7) EP300 ENCODE Project IDR conservative ChIP-seq peaks, and 8) EP300 ENCODE Project IDR optimal ChIP-seq peaks. For each definition, the following steps were taken to create positive and negative samples for the model training. For positive labels, each individual cell line DNase-I hyper-sensitive site (DHS) regions were taken and intersected with its respective definition peak file using the BEDTools command line tool (34). Any regions overlapping with promoter regions spanning 1.5 kbp upstream and 0.5 kbp downstream from each alternative transcription start site were removed (as defined in knownGene table of UCSC Genome browser, Table S16). The remaining regions were centered on the peak of the intersecting DHS sequence and extended to 1kbp, and assigned a one in the target label vector. DHS regions from all cell lines established the control set, and any overlaps with any of the defined enhancer regions were removed. Chromosomes 8 and 9 were used exclusively for testing and removed from training and validation. Chromosome 7 was used for validation, and all other autosomes were used in the training dataset. ChromHMM Strong Enhancer States and ChromHMM Weak Enhancer States were extracted from the NIH Roadmap Epigenomics project (6) (Table S10). The DHS, H3K27ac, H3K4me1, and H3K4me3 peak regions were obtained from the UCSC ENCODE database (1; 2; 3; 4; 5; 7) (Table S11, S13, S14, S15). The EP300 IDR optimal and conservative ChIP-seq peak call sets came from the ENCODE portal (Table S12) (Cherry et al. 2016, 2017, 2020) (https://www.encodeproject.org/) with the following identifiers: ENCFF087TMV, ENCFF843SYD, ENCFF583XDA, ENCFF784YVX, ENCFF054JWD, ENCFF259JWD, ENCFF821GNB, ENCFF549TYR, ENCFF476RII, ENCFF222XPQ, and ENCFF227EUK.

### Experimental raQTL Validation

RaQTL experimental data was pulled from the Open Science Framework (OSF) raQTL list (Table S17), where 14,183 single nucleotide polymorphisms (SNPs) from the HepG2 cell line and 19,237 SNPs from the K562 cell line were identified as significantly impacting the activity of regulatory regions in MPRA assays. The eight enhancer definitions and corresponding control sets were created with the same previously outlined methodology used in *Enhancer Scoring Validation* for these two cell lines.

We performed in silico mutagenesis on the position of raQTL (for both reference and alternative alleles), by calculating the change of enhancer score on a 1kbp DNA segment centered at the raQTL. These two sequences were scored by the enhancer model, and the expression:

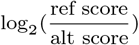

was taken to yield the delta score for each individual raQTL in the two datasets. The entire dataset of raQTLs was then separated into five equal bins based on the percentile of delta scores for each cell line (20% each). Next, the raQTL density in enhancer regions was calculated by taking the number of raQTLs in each percentile bin that overlapped with the defined enhancers divided by the total number of enhancers. For the control group, 1,000 random samples were taken where 20% of the raQTLs and a random selection of DHS regions equal to the number of enhancer regions and of equal length were taken, and this same calculation was performed. This one fifth of raQTLs was meant to standardize for the raQTL binning distribution.

These calculations were done for all delta scores and positive delta scores (SNPs that decreased enhancer score) for each of the eight enhancer definitions. This control measurement was used to calculate the fold change for each of the experimental group bins, defined as the density of raQTL SNPs in the defined enhancer regions for that given bin divided by the density of raQTL SNPs in the control set. An extensive collection of these plots can be found in Fig S4, S5, S6.

## Results

### ChromDL Architecture Discovery

Identification of this model architecture involved investigating numerous different combinations of convolutional and recurrent neural network layers, as well as pooling, normalization, and regularization alterations using random sampling and semi-supervised architecture design. The first approximately 1,100 models generated were constructed similar to DeepSEA and traditional CNN architectures like AlexNet (35) with variable convolutional filter sizes, layers of max pooling, and batch normalization. We found that these models trained with the Adam optimizer yielded prediction auROCs of about 0.918 on average, so we explored alternative methods beyond that of the original DeepSEA architecture. We then investigated if improvements could be achieved by implementing hybrid neural networks, incorporating GRUs, LSTMs, and simple RNN layers into our CNN model (about 6,000 models). With implementation, we saw an improvement in classification metrics yielding best case auROCs of approximately 0.92-0.925, and next moved to including several of these layers either of the same or different type in a single model. We found that the inclusion of multiple of these layers in the final iterations of the models as well as introducing bidirectionality within the RNN layers (about 705 models) generated the best output metrics (average auROC scores in the high 0.93 range when run for 20 epochs), with ChromDL emerging as the clear best model in terms of both median auROC (0.961) and area under the precision recall curve (auPRC; 0.402).

ChromDL exhibited an accuracy of 0.970, 0.936 and 0.864 in median auROC for ChIP-seq TFBS, DHS, and HM, respectively (Fig 2). We also observed that ChromDL outperforms DeepSEA in 95% of the 919 labels, DanQ – in 88%, and DanQ-JASPAR – in 85% (auROC; Table 1, S2; Fig S7, S8, S9) and outperforms these DL models across all categories with the single exception of DanQ-JASPAR in HM labels (Table 2, 3). We attribute the superior accuracy of ChromDL predictions to the presence of the BiGRU input layer and two BiLSTM layers, as the other network architecture elements are similar between ChromDL and the previously published models.

**Table 1.**
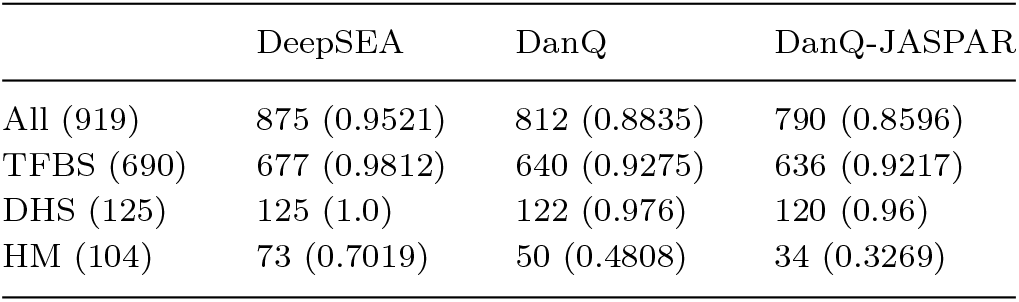
The number and proportion of labels for each of the four categories of labels (All, TFBS, DHS, and HM) that ChromDL has a higher area under the receiver operating characteristic curve (auROC) in a label-by-label comparison with each model individually.

**Table 2.**
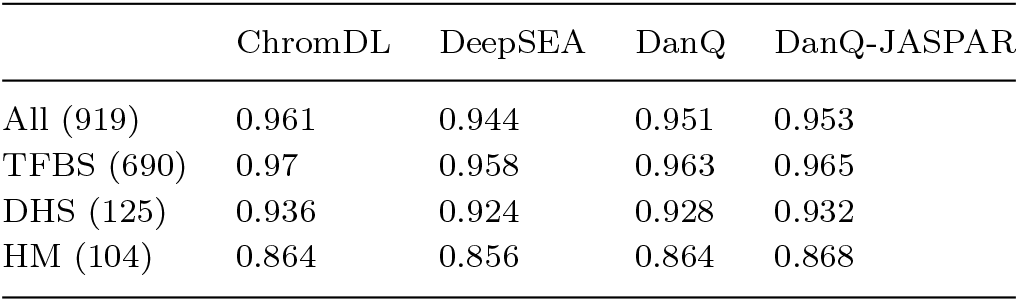
Median area under the receiver operating characteristic curve (auROC) for the four studied models for All 919 DeepSEA dataset labels, 690 TFBS labels, 125 DHS labels, and 104 HM labels.

|  | ChromDL | DeepSEA | DanQ | DanQ-JASPAR |
| --- | --- | --- | --- | --- |
| All (919) | 0.961 | 0.944 | 0.951 | 0.953 |
| TFBS (690) | 0.97 | 0.958 | 0.963 | 0.965 |
| DHS (125) | 0.936 | 0.924 | 0.928 | 0.932 |
| HM (104) | 0.864 | 0.856 | 0.864 | 0.868 |

**Table 3.**
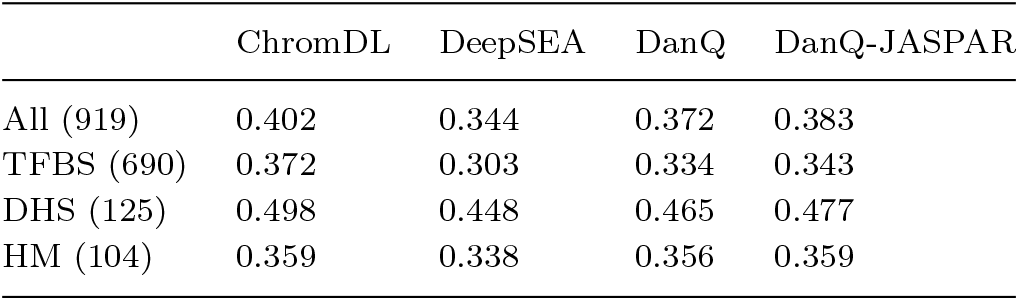
Median area under the precision recall curve (auPRC) for the four studied models for All 919 DeepSEA dataset labels, 690 TFBS labels, 125 DHS labels, and 104 HM labels.

|  | ChromDL | DeepSEA | DanQ | DanQ-JASPAR |
| --- | --- | --- | --- | --- |
| All (919) | 0.402 | 0.344 | 0.372 | 0.383 |
| TFBS (690) | 0.372 | 0.303 | 0.334 | 0.343 |
| DHS (125) | 0.498 | 0.448 | 0.465 | 0.477 |
| HM (104) | 0.359 | 0.338 | 0.356 | 0.359 |

**Fig. 2:**
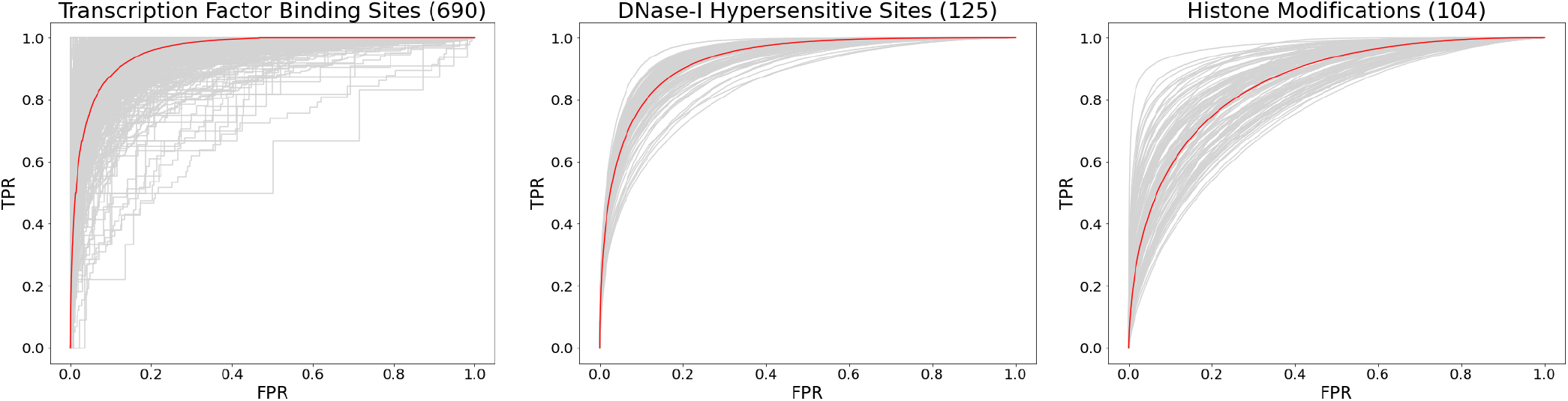
ChromDL ROC curves for TFBS, DHSs, and HMs for each available chromatin feature file across the three categories. The red line in each plot represents the average of the ROC curves across every dataset available in the trials, with each gray line representing one dataset.

To evaluate the contribution of each layer of ChromDL, we performed nine separate trials and removed one individual layer in each trial from the final architecture and evaluated the change in performance. It is important to note that while these individual layers are crucial for the model to achieve the performance metrics it does, the entire model and each layer’s variables work together to yield its final predictions. As a result, it is difficult to value one layer over another because of numerous hyperparameters that are involved and the intricacies of the individual layers. Even so, we show that the input BiGRU layer yields higher overall prediction accuracy (auROC/auPRC) as compared to a conventional CNN layer used as the input for this model (Table S3).

### Accurate Prediction of TF binding

Next, we investigated the correlation between the prediction accuracy of TF bound regions and TF ChIP-seq signal intensity. The TF ChIP-seq signal intensity reflects the propensity of TF binding, so we were interested in addressing how our model would detect the regions that had a lower binding affinity as compared to the previously developed models. We observed that ChromDL predicts a statistically significantly higher proportion of TFBS correctly in the lowest ChIP-seq signal strength bin compared to DeepSEA, DanQ, and DanQ-JASPAR (p-value *<* 1E-05; one-tailed t-test). This statistically significant improvement remains present for all incremental bins of signal strength (ranging from 0-10% to 90-100%) and across 3%, 5%, and 10% false positive rate thresholds (FPR) except for the highest two scoring bins in the 10% FPR calculations, asserting our model’s superior ability to predict TFBS at low signal strengths (Fig 3, S10, S11, Table S4, S5, S6).

**Fig. 3:**
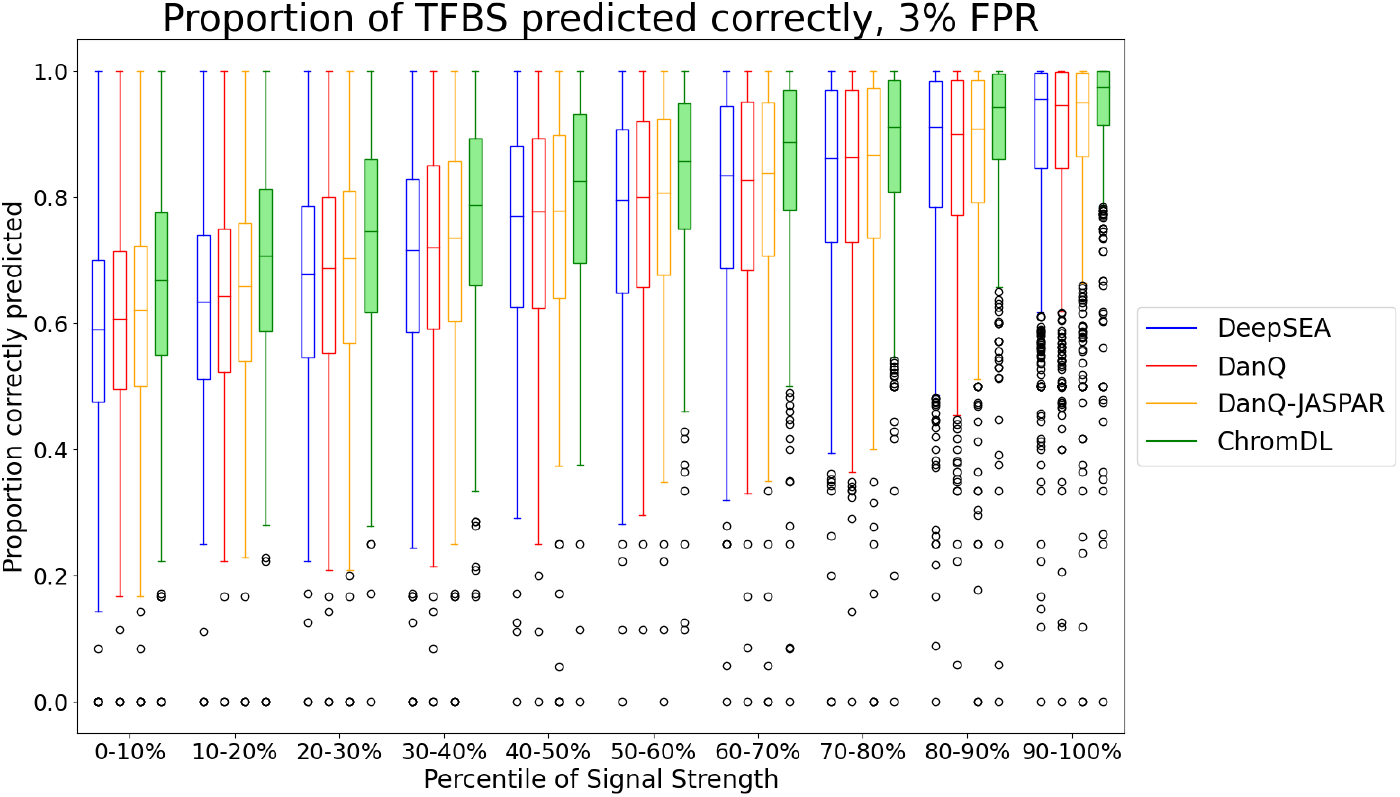
The proportion of TFBS ChIP-seq peaks in chromosomes 8 and 9 predicted correctly based on the ChIP-seq signal strength. The 3% false positive rate thresholds were calculated from the DeepSEA dataset and used to generate the plots for each of the 690 available ChIP-seq peak files.

Some of the weak signal TF ChIP-seq peaks represent sequencing errors and transient TF binding. As a result, their detection can often be difficult using predictive models due to noise associated with the corresponding signal levels as compared to the strong signal TF ChIP-seq peaks. It is therefore significant that our model can detect and predict these weak signal TFBS regions with higher accuracy as compared to previously developed models. This higher sensitivity could allow detection of sequencing and mapping errors in existing datasets, as well as open the possibility of detecting transiently bound regions with improved accuracy.

To address ChromDL’s ability of predicting TF binding motifs, we utilized Protein Binding Microarray (PBM) profiling of in vitro DNA-binding of transcription factors (36). A total of sixteen motifs that were found in both the DeepSEA TFBS dataset and the PBM published database were examined in this investigation. For these motifs, the average number of peaks was 1,276.5, and the average percentage of these peaks predicted correctly was 81% for ChromDL, 75% for DeepSEA, 75% for DanQ, and 77% for DanQ-JASPAR. Eight of the 16 motifs were not visually discernible with the PBM PWMs and resulted in insignificant TOMTOM alignment mapping scores, so they were discarded. The other eight motifs, all c-Fos/c-Jun, were detected as a prominent motif in each of the four comparison models (ChromDL, DeepSEA, DanQ, DanQ-JASPAR). For seven of the eight motifs, all four models predicted the PBM motif as the most significant motif, and one motif, HepG2 c-Jun, was the second most significant in all four models. This motif was then extracted for each model and compared with the target PBM motif to calculate alignment scores. In these two investigations, ChromDL had the most significant alignment score (lowest p-value) in four out of the eight motifs from HOMER generated motifs (Table S7), and the most significant alignment score in four out of eight motifs from MEME-ChIP generated motifs (Table S8). We found through this investigation that ChromDL not only has the ability to map specific base pairs correctly, but also is able to maintain high sensitivity to the weight of the correct individual base pair in TFBS PWM description more in-line with PBM data. This sensitivity can be visualized in Fig 4.

**Fig. 4:**
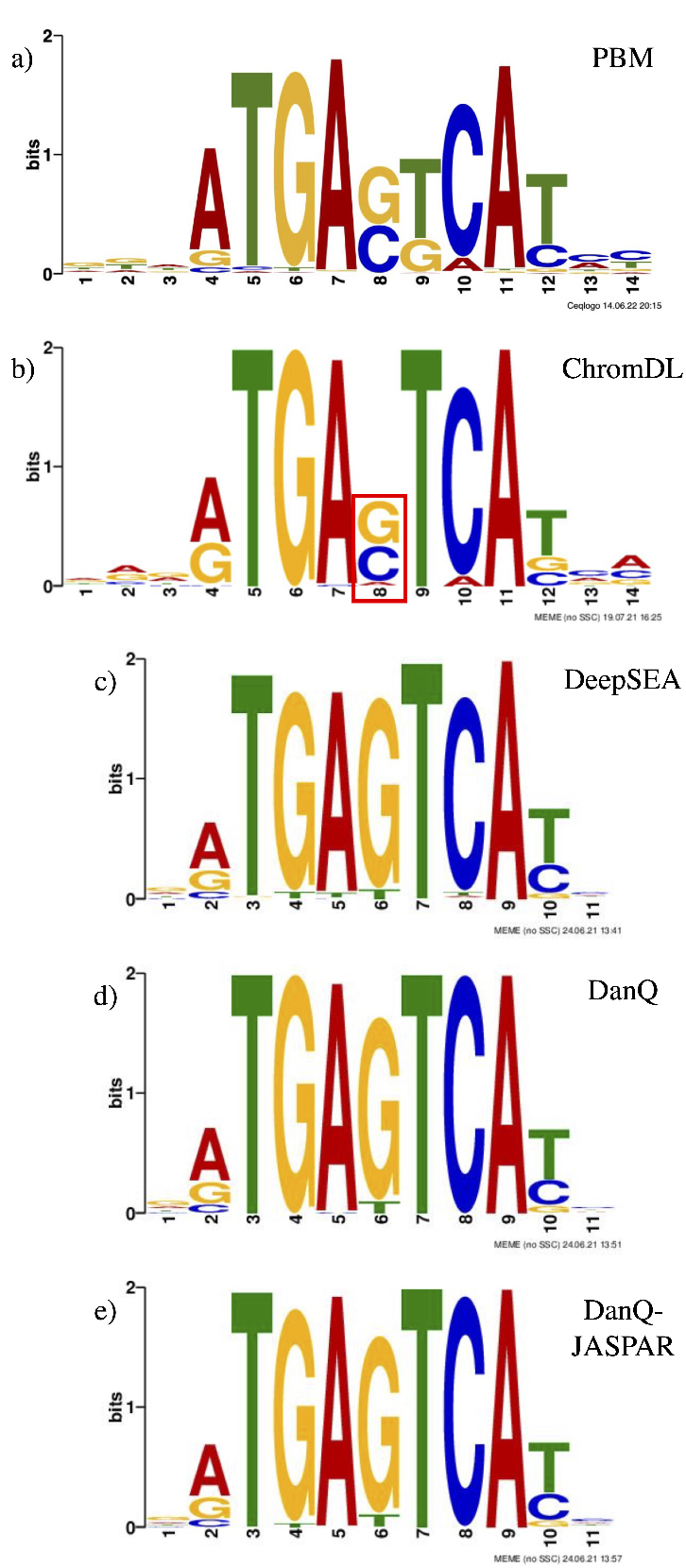
Motif analysis of Jun/Fos gene using PBM and MEME Suite. (a) primary motif logo in c-Jun/c-Fos protein in Homo Sapiens reported by the UniProbe PBM study (30; 33). The other logos correspond to the HepG2 c-Jun TFBS ChIP-seq PWMs mapped for (b) ChromDL, (c) DeepSEA, (d) DanQ, and (e) DanQ-JASPAR. Highlighted is the base pair at position 8 in the ChromDL PWM, illustrating the higher sensitivity at this position leading to a closer match with the PBM PWM.

### ChromDL-based Enhancer Predictor

Next, we used the TREDNet DL second-phase classifier CNN architecture with the output of the ChromDL model serving as input to compute an enhancer score of an arbitrary input DNA region. The two-stage model yielded respectable performance, with an average auROC of 0.89 across eight enhancer definitions (Fig S12, Table S18). We took a particular interest in the HepG2 and K562 cell lines for subsequent analysis and obtained high auROC (average 0.967 and 0.949) and auPRC metrics (average 0.313 and 0.352) in the EP300 enhancer trials as compared to other definitions (Fig S13, S14).

When we examine the trends across all available cell lines and the eight enhancer definitions, we notice several key differences. We find that the lowest scoring auPRC enhancer group are the regions centered on overlapping H3K27ac and H3K4me1 peaks without H3K4me3 peak overlaps, with an average auPRC of 0.28. Closely behind it is the ChromHMM Strong Enhancer State definition with an average auPRC of 0.33. We attribute this poor relationship between the precision and recall to the enhancer definitions, as these two groups in particular have sequence features as compared to the other groups which lead them to be previously termed “stronger” enhancers. On the other hand, the “weaker” enhancer groups, which have been demonstrated to lead to elevated enhancer activity (37), resulted in more accurate ChromDL classification according to the auPRC metric. It has also been experimentally demonstrated that weaker regions with fewer H3K27ac peaks and more H3K4me1 peaks lead to an increased enhancer activity, and we would thus expect it to lead to a better auPRC metric. Indeed, we find this to be true in our overlapping H3K27ac and H3K4me1 peak enhancers with an average auPRC of 0.40, and our H3K4me1 peak enhancers with an average auPRC of 0.55. These results are encouraging, since the high auROC (*>*0.84 average across all definitions) suggest that ChromDL is able to provide enough contextual information to this enhancer model across multiple cell lines.

### Reporter assay QTL validation of ChromDL Regulatory Variant predictions

After including the TREDNet second-phase enhancer classifier for the prediction of enhancer sequences using the output context from ChromDL as input, we decided to address if the enhancer scores are sensitive to point mutations in the input enhancer regions. We found that indeed, this two-stage model can be directly used to quantify the impact of DNA sequence mutations on enhancer activity by computing the difference in enhancer scores corresponding to two different alleles of the mutated nucleotide.

We performed an in silico mutagenesis of every nucleotide of HepG2 EP300 enhancers (optimal IDR) using the ChromDL enhancer model and observed that 52.8% of enhancer mutations show a detrimental impact on enhancer activity with 2.2% of enhancer mutations being capable of enhancer deactivation (defined as at least a 2-fold enhancer score decrease), which is in line with previously reported results of deleterious enhancer mutations (38; 39; 40; 41). Additionally, 47.2% of enhancer mutations are strengthening enhancer activity and 0.65% of mutations increase the ChromDL enhancer score over 2-fold. These results extend the scope of the previously reported enhancer gain mutations for a single ZRS enhancer to the whole-genome scale and identify a small set of key mutations with the most pronounced impact on enhancer activity (42).

To validate the accuracy of ChromDL regulatory variant predictions, we used the results of experimental characterization regulatory effects at Single Nucleotide Polymorphism (SNP) positions in the human genome using the raQTL technique (43), which mapped 14,183 and 19,237 regulatory SNP mutations in HepG2 and K562 cell lines, respectively.

For each raQTL, two sequences were created and fed into the pre-trained enhancer model attached to ChromDL, a 1kbp sequence centered at the wild type allele and a 1kbp sequence centered at the same allele with the mutant base pair in its place. Those raQTLs with the greatest delta scores (corresponding to the greatest predicted impact on enhancer activity) were found to be the most enriched in enhancer regions (Fig 5). These results indicate that the ChromDL enhancer classifier can be applied for the accurate prediction of regulatory mutations and the degree of enhancer modulation by these mutations. Our model has been experimentally validated through 1) The trend of increasing magnitude raQTL score correlating with higher density in enhancer sequences and 2) the proportion of enhancers that were deactivated (2-fold score decrease) in the mutagenesis trials aligning with previously published results.

**Fig. 5:**
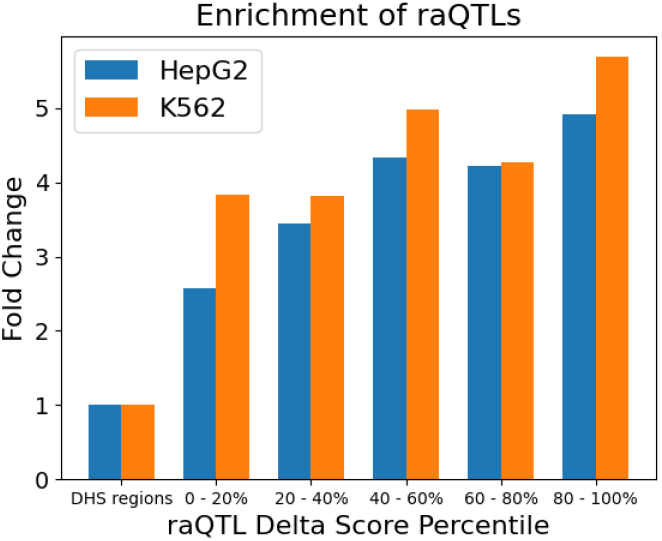
raQTL enrichment binned by DL delta score.

## Discussion

By testing thousands of various DL architectures, we have built an accurate DL predictor of regulatory activity in the human genome named ChromDL. ChromDL improves the accuracy of prediction of regulatory features as compared to DeepSEA, DanQ, and DanQ-JASPAR. The optimal ChromDL model highlights the benefits of RNN layers for prediction of genomic features as compared to the traditional CNN architecture. We also attribute the presence of the two BiLSTM layers and a BiGRU layer to the improved predictive ability of the developed classifier.

ChromDL demonstrates an improved prediction accuracy in detecting low-affinity TF binding and TF binding motif identification when compared to DeepSEA and DanQ. This was demonstrated through PBMs, which are used to measure the protein-DNA binding affinity in several orders of magnitude, meaning that they are a very powerful tool in the comparison of DNA-binding specificity. It is significant as a result that ChromDL has the potential to predict these key base pairs with more accuracy as compared to DeepSEA and DanQ models.

In addition, we find that ChromDL can be utilized with an enhancer model such as the TREDNet second-phase classifier for the prediction of enhancer sequences that are sensitive to regulatory region mutations as validated through experimental raQTL point mutations. This illustrates the possibility of very effective enhancer classifiers using ChromDL’s output context and deep learning neural networks for the accurate prediction of regulatory mutations and to quantify the enhancer modulation of these mutations.

Further updates to this model could investigate lowering the training speed to analyze model iterations more quickly, initializing the model with motifs before training to see if scores improve, and training the model on outside datasets as they become available to make the model more versatile for the prediction of regulatory features. We expect to see further hybrid architectures of varying size, speed, and composition as effective in exploring human regulatory DNA using similar exploration techniques with the availability of faster GPUs and more sophisticated layer implementations.

We have built what we believe to be the most accurate regulatory classifier to date in ChromDL, with the ability to predict TFBSs and DHSs with performance metrics beyond that of all studied predecessors. We believe ChromDL’s sensitivity to low affinity TF binding, it’s flexibility with secondary classifiers while retaining strong performance metrics, and it’s basepair sensitivity to point mutations make it a powerful tool to aid in the development of new technologies and assist in further novel discoveries in the field of epigenetics.

## Supporting information

DeepSEA Data Sources

Enhancer Scoring Metrics

Experimental Validation Data Sources

## Acknowledgments

This work utilized the computational resources of the NIH HPC Biowulf cluster. (http://hpc.nih.gov). This research was supported by the Intramural Research Program of the National Library of Medicine, National Institutes of Health.

## Notes

https://github.com/chrishil1/ChromDL

### Competing Interest Statement

The authors have declared no competing interest.

