## Supplemental Materials for "ChromDL: A Next-Generation Regulatory DNA Classifier"

#### Running title

ChromDL

#### Supplemental Materials

**ChromDL Architecture (Input shape = (1000, 4))**:

1. Bidirectional GRU layer (128 units. Tanh activation. Output shape = (1000, 256))
2. Separable Convolution layer (750 filters. Window size: 16. Step size: 1. ReLU activation. Output shape = (985, 750))
3. Convolution layer (360 filters. Window size: 8. Step size: 1. ReLU activation. Output shape = (978, 360))
4. Maximum Pooling layer (Window size: 4. Step size: 4. Output shape = (244, 360))
5. Bidirectional LSTM layer (128 units. Tanh activation. Output shape = (244, 256))
6. Batch Normalization layer (Momentum: 0.99. Epsilon: 0.001. Output shape = (244, 256))
7. Average Pooling layer (Window size: 8. Step size: 8. Output shape = (30, 256))
8. Bidirectional LSTM layer (128 units. Tanh activation. Output shape = (30, 256))
9. Flattening layer (Output Shape = (7680))
10. Dense layer (919 units. Sigmoid activation. Output shape = (919))

**Regularization Parameters:**:

- L1 sparsity (layers 2 and 3): 1E-08
- L2 regularization (layers 2 and 3): 5E-07
- Dropout proportion (layer 5): 20%

**TREDNet-based Enhancer Architecture (Input shape = (919, 1))**:

1. Convolution layer (64 filters. Window size: 9. Step size: 1. ReLU activation. Output shape = (911, 64))
2. Batch Normalization layer (Momentum: 0.99. Epsilon: 0.001. Output shape = (911, 64))
3. Leaky ReLU layer ($\alpha$: 0.0. Output shape = (911, 64))
4. Maximum Pooling layer (Window size: 9. Step size: 3. Output shape = (301, 64))
5. Convolution layer (128 filters. Window size: 4. Step size: 1. ReLU activation. Output shape = (298, 128))
6. Leaky ReLU layer ($\alpha$: 0.0. Output shape = (298, 128))
7. Maximum Pooling layer (Window size: 4. Step size: 2. Output shape = (148, 128))
8. Convolution layer (256 filters. Window size: 4. Step size: 1. ReLU activation. Output shape = (145, 256))
9. Maximum Pooling layer (Window size: 4. Step size: 3. Output shape = (48, 256))
10. Flattening layer (Output Shape = (12288))
11. Dense layer (180 units. ReLU activation. Output shape = (180))
12. Dense layer (1 unit. Sigmoid activation. Output shape = (1))

**Regularization Parameters:**:

- Dropout proportion: layer 1: 20%, layer 5: 20%, layer 8: 50%
- L1 sparsity (layers 1, 5, and 8): 1E-04
- L2 regularization (layers 1, 5, and 8): 1E-03
- Max kernel normalization (layers 1, 5, and 8): 1

### Model Discovery

We took an approach to identify promising models from their auROC scores after a single epoch under standardized training conditions (Adam optimizer, batch size of 500). Permutations of models were strategically generated for comparison against DeepSEA using the layers available in the Tensorflow library [1, 2]. Several constraints were applied in model generation, including requiring at least one convolutional layer and a number of pooling layers relative to the size of the model. When running trials on the GPU, any models with incompatibility, memory errors, or time estimations exceeding ten hours per epoch were discarded. Thousands of generated models were trialed with the highest scoring one epoch models isolated and run for 100 epochs or until validation loss plateaued, and were then evaluated for auROC/auPRC scores. This led to a handful of models scoring at or above the published values of the primary comparison models.

The promising models were then trialed across all available tensorflow optimizers and their respective implementations. The Adadelta, Adagrad, Adam, Adamax, Ftrl, Nadam, RMSprop, and SGD optimizers were all run on the top model for 50 epochs to gauge performance [1, 2]. From these trials, the Adam optimizer was ultimately reaffirmed for this model based on auROC/auPRC scores. A summary of the ChromDL architecture testing metrics when trained with these different optimizers can be found in Table S1.

The final step in fine-tuning these models was individual layer insertions and deletions as well as hyperparameter optimizations to improve auROC scoring. These alterations ranged from including or excluding convolutional, pooling, normalization, and activation layers, changing unidirectional layers to bidirectional and vice versa, and adding in GRU, LSTM, and SimpleRNN layers. We also adjusted the number of neurons, filter sizes, window sizes, pooling sizes, batch sizes, and L1/L2 terms in the layers. These altered models were run for 20 epochs, and the highest scoring trials were then run until the validation loss plateaued for 20 epochs, capped at 100 epochs. Improvements in final auROC/auPRC scores were observed from this methodology, and this final model achieved the highest scoring results of this process.

### ChromDL Layer Specifications

Whereas DeepSEA and DanQ both process the one-hot encoded DNA sequence input data with a convolutional layer, ChromDL instead uses a bidirectional gated recurrent unit (BiGRU) as the input layer [3, 4]. Briefly, the BiGRU layer is made up of two parts, the GRU layer itself and the bidirectional wrapper. The GRU is a variation of a simple RNN, where a fully connected layer is created which feeds the output back into the input layer, with the output being used as the state. A GRU attempts to improve upon this design and eliminate the vanishing gradient problem with two additional units: an update gate and a reset gate [5]. This layer can be expressed as:

$$r_j = \sigma([W_r x]_j + [U_r h_{t-1}]_j)$$
$$z_j = \sigma([W_z x]_j + [U_z h_{t-1}]_j)$$
$$h_j^t = z_j h_j^{t-1} + (1 - z_j)\tilde{h}_j^t$$
$$\tilde{h}_j^t = \phi([Wx]_j + [U(r \odot h_{t-1})]_j)$$

Where $x$ is the input vector, $r_j$ is the reset gate, $z_j$ is the update gate, $h_j$ is the hidden unit, $\tilde{h}_j$ is the new hidden state, $h_{t-1}$ is the previous hidden state, $[...]_j$ denotes the jth element in the vector, $W$ and $U$ are learned weight matrices, $\sigma$ is the logistic sigmoidal function, $\phi$ is the hyperbolic tangent, and $\odot$ is an element-wise multiplication [5].

This sigmoidal activation function acting on the update and reset gates is expressed as:

$$\text{Sigmoid}(x) = \frac{1}{1 + e^{-x}}$$

And the hyperbolic tangent activation function acting on the hidden state is expressed as:

$$\tanh(x) = \frac{e^x - e^{-x}}{e^x + e^{-x}}$$

The update gate determines the amount of information from previous input reads that is passed to the next state, and the reset gate determines how much of the past information is forgotten. Unlike a traditional RNN or an LSTM, the GRU does not contain an output gate and instead has this update gate which combines the input from the current read and the reset gate.

In a bidirectional layer, the sequences are processed from beginning to end and from end to beginning simultaneously, with two identical hidden layers accounting for each direction. By doing so, the layer can learn from past, present, and future traversal of input in adjusting the weights applied before passing the output to the next layer [6]. In a BiGRU layer, we have all the same layer variables as before, with the only difference between the two hidden layers being the starting point of the sequence read. The next layer receives the functional transformations applied from both of the hidden layers simultaneously.

Following the BiGRU input layer, the sequences are fed into a separable convolutional layer. This layer differs slightly in implementation from the traditional convolutional layers found in DeepSEA and DanQ [3, 4], where the output is calculated through a one dimensional spatial convolution with a number of filters and window size [4, 7]. In a traditional CNN, each of these filters is a weight matrix similar in concept to the neurons of other layers, with scores calculated as a sliding window with a fixed step size moves across the sequence and extracts features of the input [7]. Mathematically this can be expressed as:

$$\text{Conv}(X)_{ik} = \text{ReLU}(\sum_{m=0}^{M-1} \sum_{n=0}^{N-1} W_{mn}^k X_{i+m,n})$$

Where $X$ is the input, $i$ is the index of the ouput position, and $k$ is the index of the filter. Each filter is represented as $W^k$ and is a $M$ x $N$ weight matrix where $M$ is the window size and $N$ is the number of input channels [4, 7]. Typically after convolution, the rectified linear unit (ReLU) activation function is applied to the output before it is passed to the next layer. This activation function sets all negative input values to zero and not affect any value greater than zero [8, 9], and can be expressed as:

$$\text{ReLU}(x) = \begin{cases} x, & \text{if } x > 0 \\ 0, & x \leq 0 \end{cases}$$

Before the traditional convolutional layer in our model is a separable convolutional layer, which as implemented in the Tensorflow library first performs a depthwise convolution that acts separately on channels, followed by a pointwise 1x1 convolution that mixes the channels together [2, 7]. These separable convolutions are thought to reduce the number of parameters while increasing efficiency of the resulting feature map representation [7]. Our model takes the output after rectified linear unit (ReLU) activation and passes it through a traditional convolutional layer, also with ReLU activation.

Following activation, the input is maximally pooled. This pooling layer calculates the maximum value in a window with an equal step size to reduce the size of the output while still maintaining the key spatial features and minimizing memory overhead. This max pooling step is expressed as:

$$\text{MaxPool}(X)_{ik} = \max(X_{(iM,k)}, X_{(iM+1,k)}, ..., X_{(iM+M-1,k)})$$

Where $X$ is the input to the layer, $i$ is the index of the output layer, $k$ is the index of the inputted layer (in this case the index of the filter), and $M$ is the pooling window size [4, 8, 9].

We then have the first of two BiLSTM layers, which contains both an LSTM layer and a bidirectional wrapper that creates a forward and reverse hidden layer. Briefly, the expression for the LSTM layer is as follows:

$$\begin{aligned} f_t &= \sigma(W_{fx}x_t + W_{fh}h_{t-1} + b_f) \\ i_t &= \sigma(W_{ix}x_t + W_{ih}h_{t-1} + b_i) \\ o_t &= \sigma(W_{ox}x_t + W_{oh}h_{t-1} + b_o) \\ \tilde{c}_t &= \phi(W_{cx}x_t + W_{ch}h_{t-1} + b_c) \\ c_t &= f_t c_{t-1} + i_t \tilde{c}_t \\ h_t &= o_t \phi(c_t) \end{aligned}$$

In this definition, $i_t$ is the input variable, $c_t$ is the cell variable, $f_t$ is the forget variable that controls how much information of $c_{t-1}$ is remembered, $\tilde{c}_t$ is the new information to save, and $o_t$ is the output variable that controls how much of $c_t$ is saved to $h_t$, the hidden variable [10, 11]. $W$ is the weight matrices of these different variables, $b$ is the bias vector for each of the four variables, $\sigma$ is the sigmoidal function, and $\phi$ is the hyperbolic tangent [10, 11]. A bidirectional LSTM has two of these units as hidden layers, one that moves in the forward direction and one in the reverse direction, simultaneously feeding their results into the next sequential layer.

Next we have a dropout layer, which will randomly set a proportion of the neurons (in this case in the previous BiLSTM layer) to zero to prevent overfitting of the input data [12]. This will occur over each step of the training with a different random proportion of the neurons, emphasizing the importance of training over many epochs.

A batch normalization layer is next, which applies a transformation that attempts to keep the mean output close to zero and the mean standard deviation of the output close to one [13]. The mathematical expression that reflects this is as follows:

$$y_i = \frac{\gamma(x_i - \mu_B)}{\sqrt{\sigma_B^2 + \epsilon}} + \beta$$

Where $x$ is the input to the normalization layer, $y$ is the output of the transformation, $\mu_B$ is the mini-batch B mean. $\sigma_B$ is the mini-batch $B$ standard deviation, and $\epsilon$ is a constant for numerical stability. $\beta$ and $\gamma$ are learned through training and apply a channel-wise affine transformation [13].

Following the normalization layer is an average pooling layer. This functions the same way as the max pooling layer, but instead of taking the maximum value in the sliding window, the average value is taken. This is expressed as:

$$\text{AveragePool}(X)_{ik} = \text{avg}(X_{(iM,k)}, X_{(iM+1,k)}, ..., X_{(iM+M-1,k)})$$

Where $X$ is the input to the layer, $i$ is the index of the output layer, $k$ is the index of the inputted layer, and $M$ is the pooling size window. We tried numerous combinations of average pooling and maximum pooling using various window sizes and ultimately found that this maximum pooling followed by average pooling yielded the highest prediction scores. Following this, we have another BiLSTM layer before the input is passed into a flattening layer, where all of the input is flattened to one dimension in order to be processed in the final output layer.

This final output layer is a dense layer of neurons that serves as a series of weight matrices applied to the inputs to get the final output values for each of the 919 targets. We use a sigmoidal output layer as an activation function, which transforms the data and combined with the weight matrices yields the final output for each of these regulatory features. Analogous to DeepSEA, all of the neurons in the output layer share the same set of input from the previous layer, so sharing of the predictive sequence features can take place across all of the 919 labels [4].

#### auPRC Investigation

In addition to the area under the receiver operating characteristic (auROC) comparisons, we conducted comparisons using the area under the precision recall curve (auPRC) metric. We found that ChromDL had respectable auPRC median scores for TFBS (median auPRC = 0.372), DHS regions (median auPRC = 0.498) and HMs (median auPRC = 0.355), outperforming the three comparison models for TFBS and DHS labels and DeepSEA and DanQ for HM labels (see Manuscript Table 3).

When examining the auPRC scores label by label, we find that our model improves prediction performance as compared to DeepSEA in HMs (outperforming DeepSEA in 81% of labels), in TFBS (in 98% of labels) and DHS sites (in 100% of labels). With the exception of HMs, we find that our model outperforms both DanQ and DanQ-JASPAR in TFBS labels (outperforming DanQ in 84% of labels, DanQ-JASPAR in 79%) and DHS labels (outperforming DanQ in 99% of labels, DanQ-JASPAR in 97%) (Table S2).

#### Enhancer Scoring LeakyReLU

Included in the TREDNet second-phase enhancer classifier is a LeakyReLU activation layer [14]. This activation layer is a variant of the ReLU activation function that allows a small non-zero gradient $\alpha$ curve when negative values occur in an attempt to solve the dying ReLU problem [15]. Briefly, this is a well documented issue where neurons that always output negative values and are subsequently set to zero by the ReLU function are essentially "dead" and do nothing to contribute to the network. The LeakyReLU activation can be mathematically expressed then as:

$$\text{LeakyReLU}(x) = \begin{cases} x, & \text{if } x > 0 \\ \alpha x, & x \leq 0 \end{cases}$$

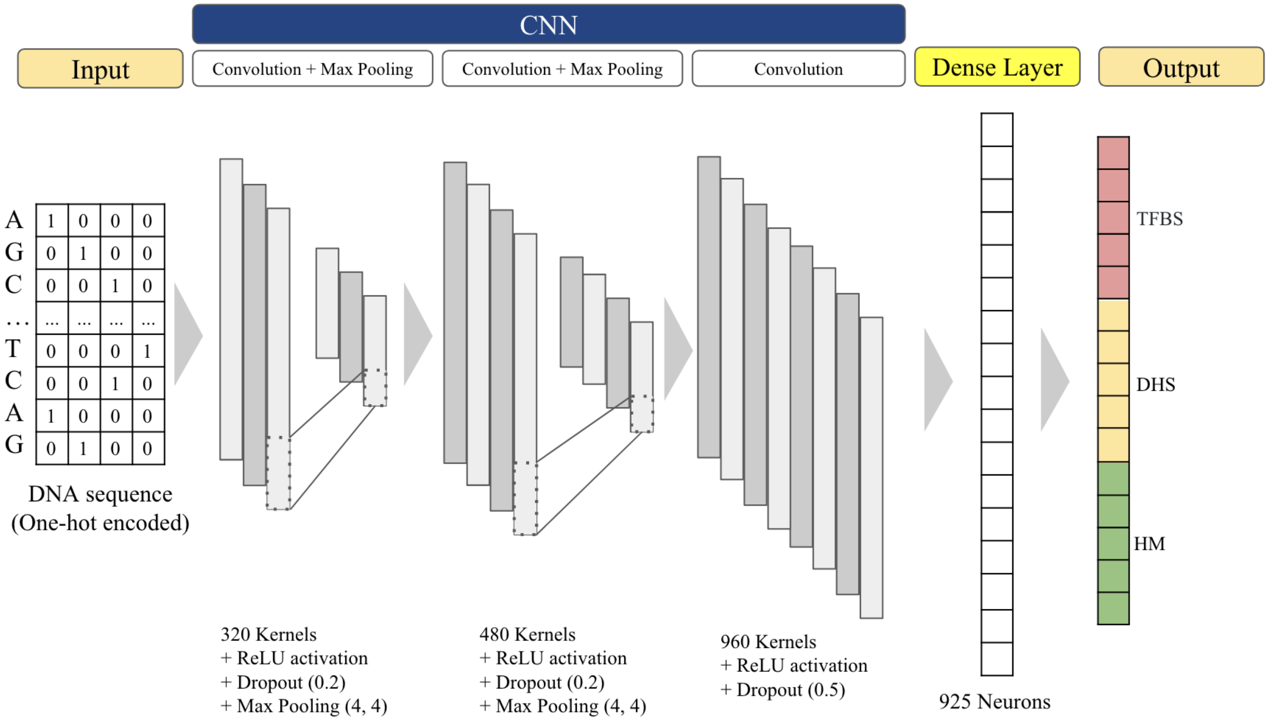


Figure 1: Visual representation of DeepSEA's DL architecture. The input sequences are fed into the model as their one-hot encoding, and processed by each Convolutional Neural Network (CNN) layer before prediction scores are calculated for the 919 chromatin features.

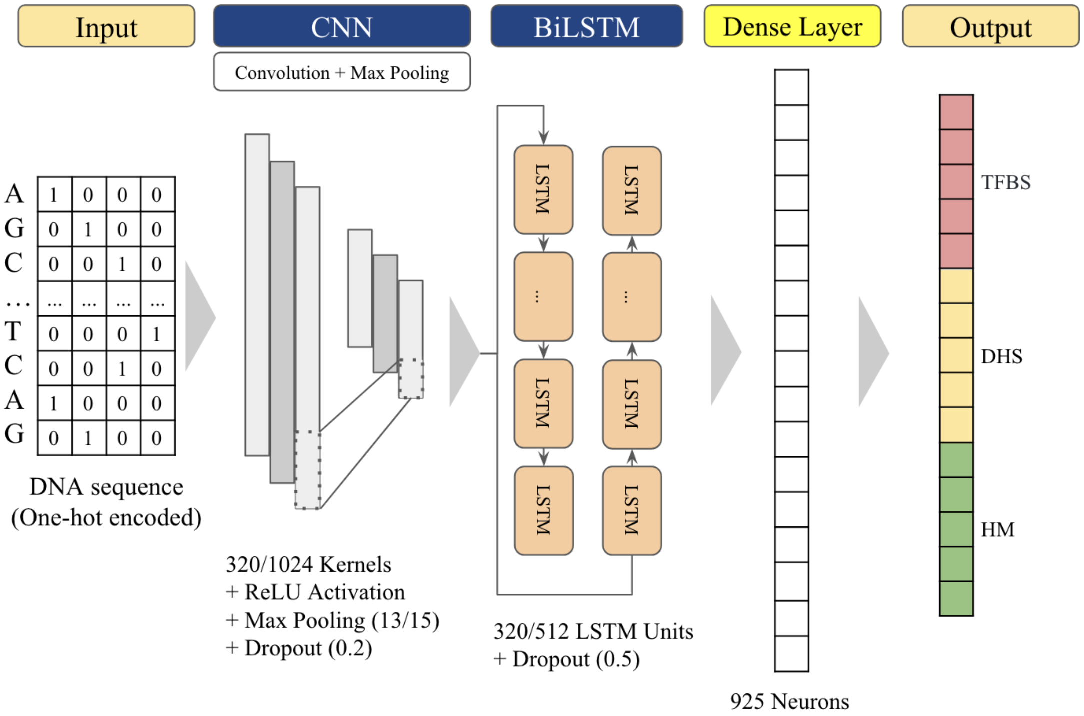


Figure 2: Visual representation of DanQ DL architecture, with backslashes denoting the parameter for the DanQ/DanQ-JASPAR model. The input sequences are fed into the model as their one-hot encoding, and processed first by a Convolutional Neural Network (CNN), followed by a Long Short-Term Memory recurrent neural network before prediction scores are calculated for the 919 chromatin features.

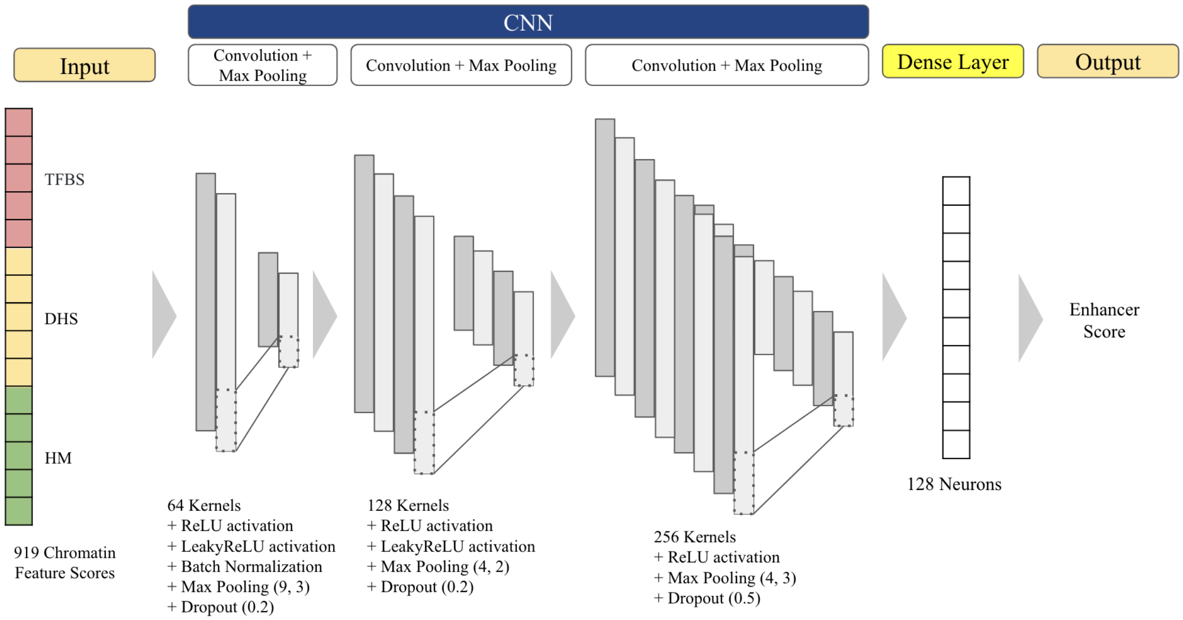


Figure 3: Visual representation of the adapted TREDNet enhancer classifier built onto the output of ChromDL's DL architecture. This classifier takes the outputted prediction scores calculated by ChromDL as input sequences and processes them through three Convolutional Neural Network (CNN) layers before one enhancer prediction score is produced.

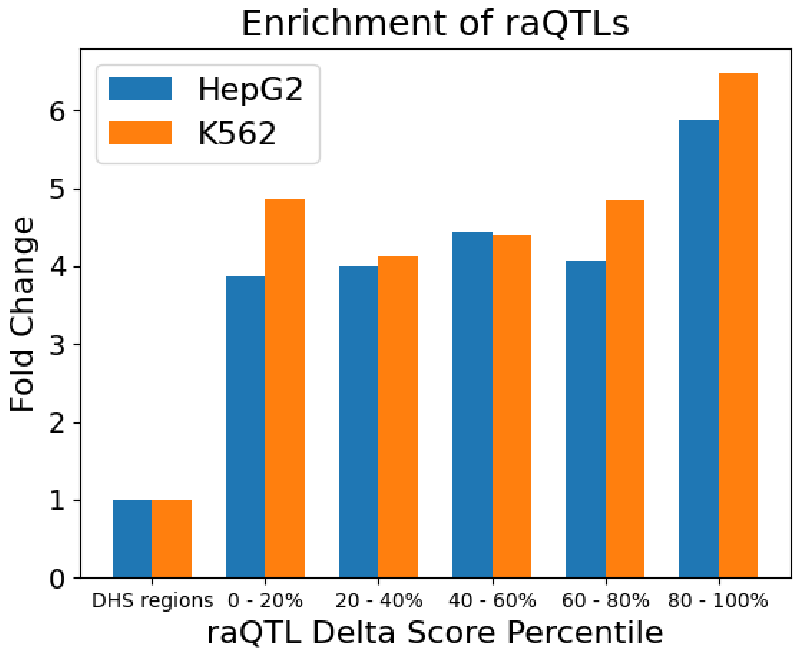


Figure 4: raQTL enrichment binned by DL delta score looking only at the positive raQTLs, or those whose mutation caused the enhancer prediction score to decrease.

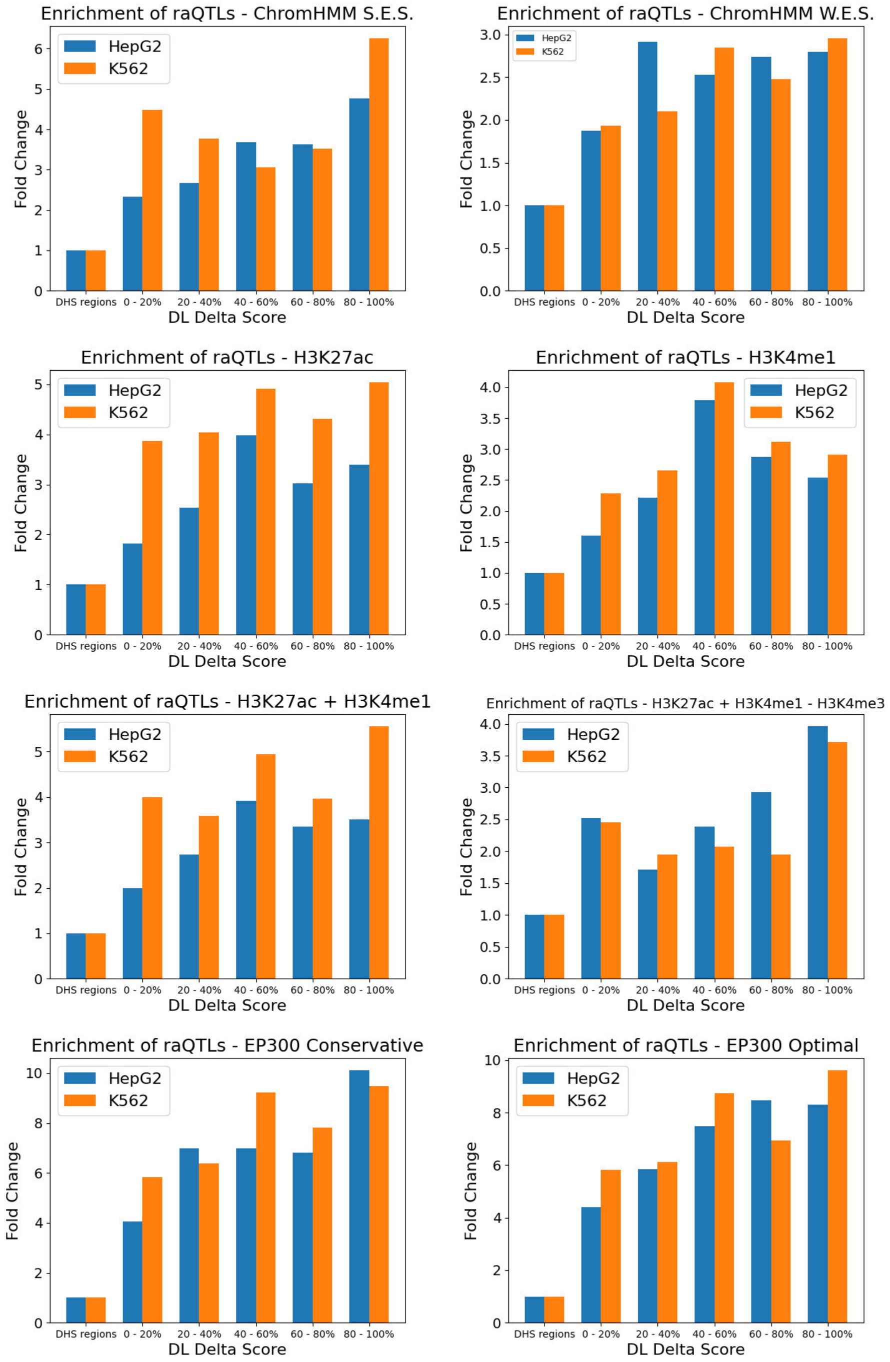


Figure 5: raQTL enrichment binned by DL delta score in the eight defined enhancer regions.

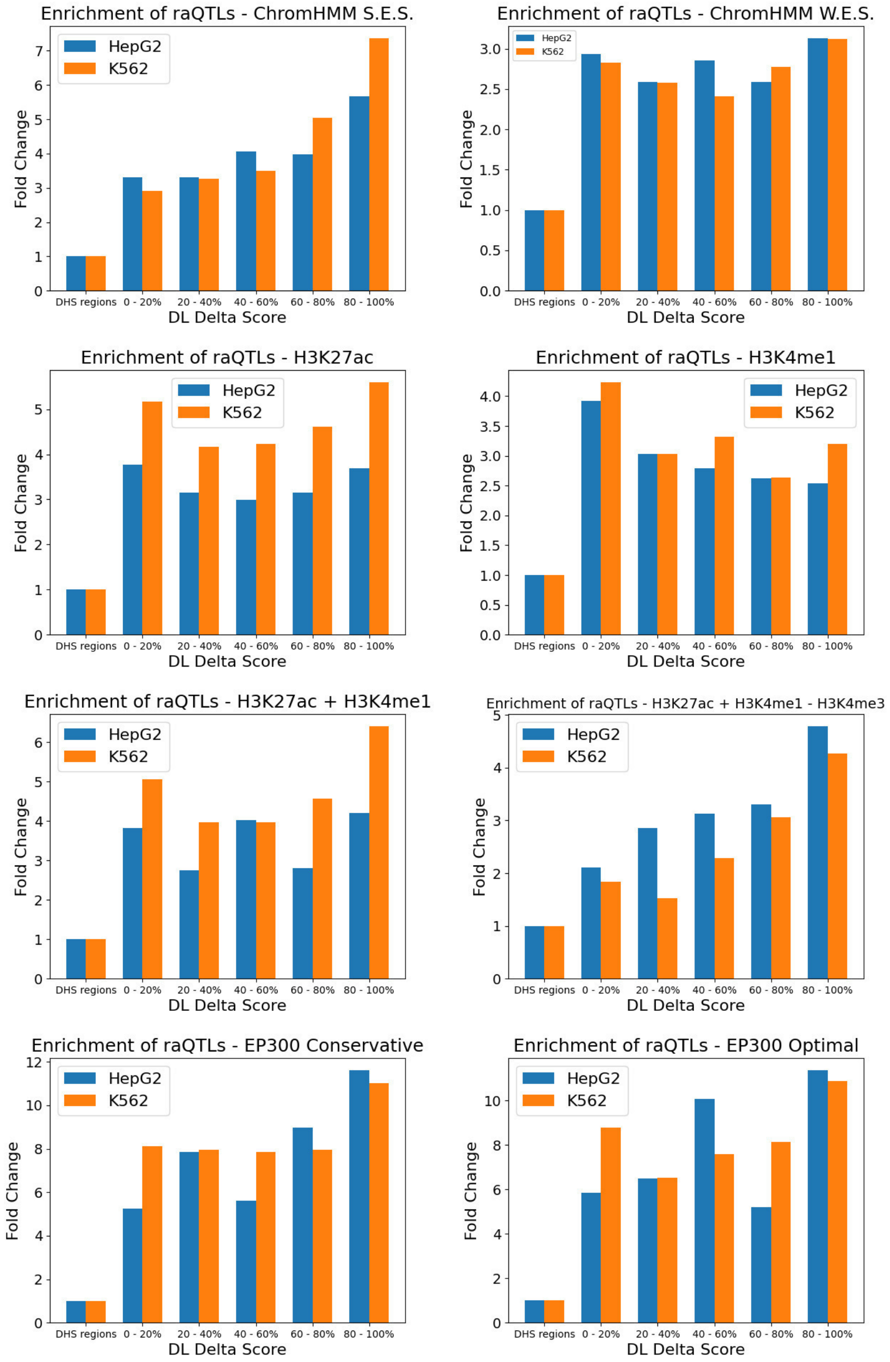


Figure 6: raQTL enrichment binned by DL delta score in the eight defined enhancer regions looking only at the positive raQTLs, or those whose mutation caused the enhancer prediction score to decrease.

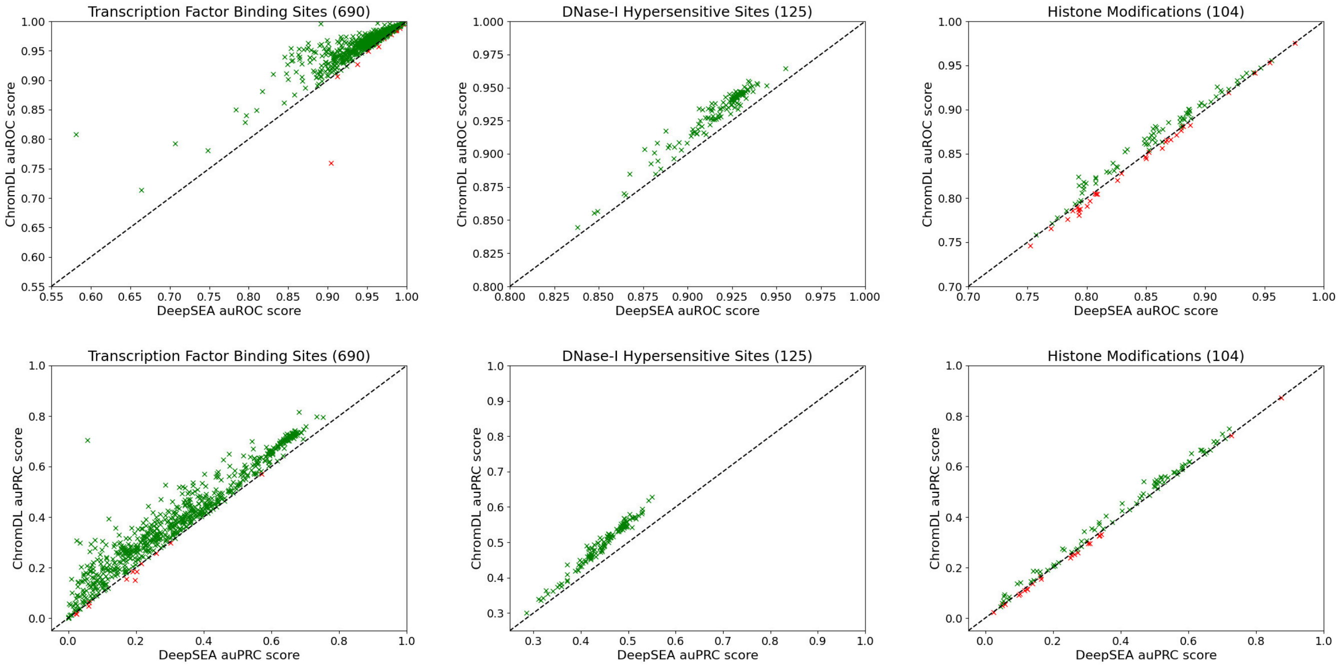


Figure 7: Label performance of ChromDL compared to DeepSEA. For a given point, x is the auROC/auPRC score from DeepSEA, and y is the auROC/auPRC score from ChromDL. Green indicates that ChromDL scored higher, and red indicates DeepSEA scored higher.

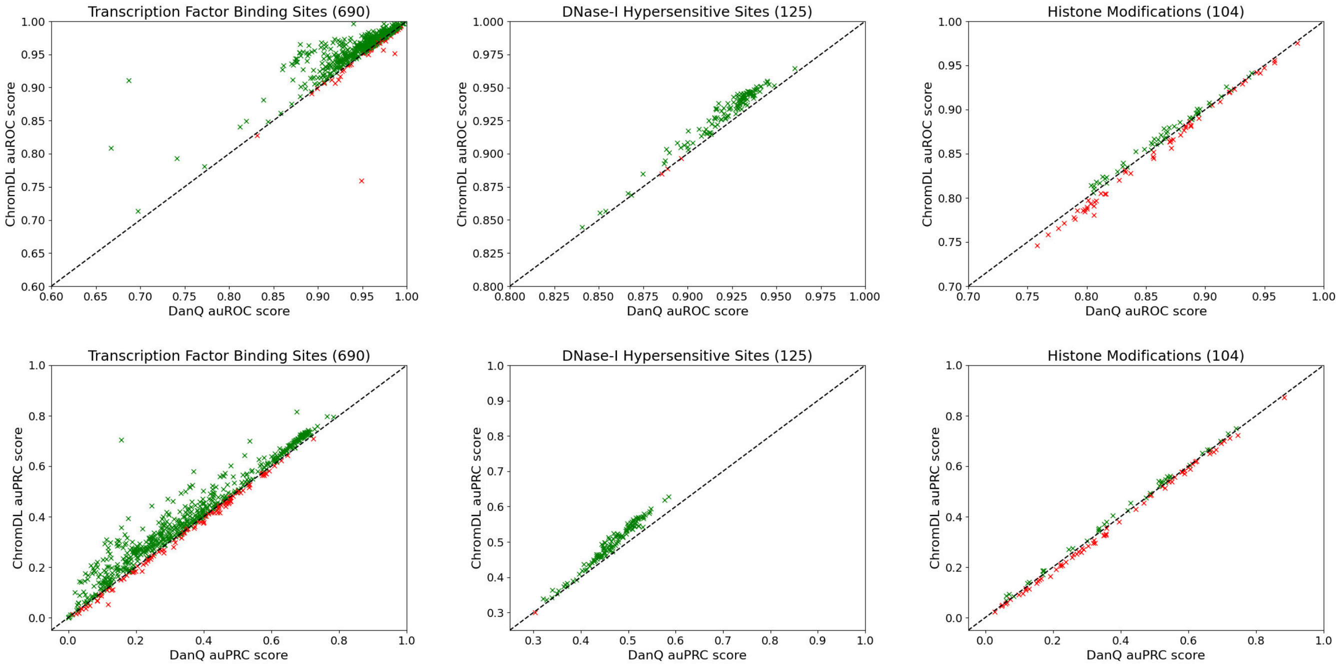


Figure 8: Label performance of ChromDL compared to DanQ. For a given point, x is the auROC/auPRC score from DanQ, and y is the auROC/auPRC score from ChromDL. Green indicates that ChromDL scored higher, and red indicates DanQ scored higher.

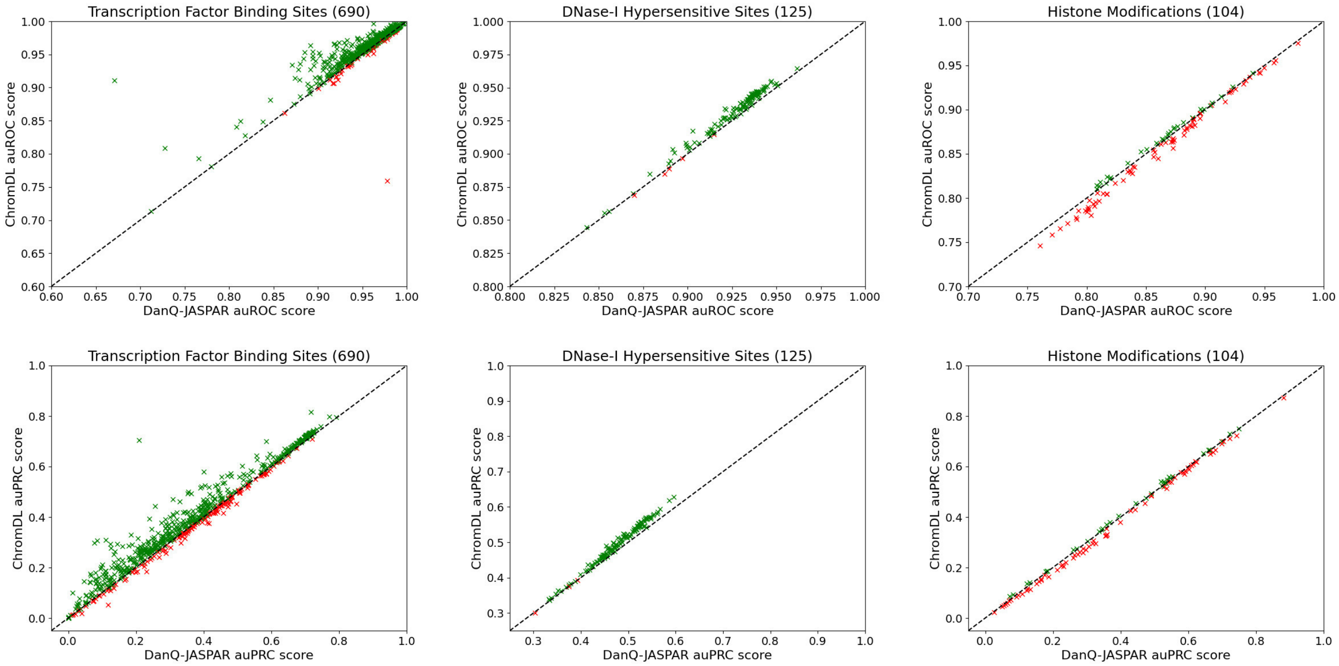


Figure 9: Label performance of ChromDL compared to DanQ-JASPAR. For a given point, x is the auROC/auPRC score from DanQ-JASPAR, and y is the auROC/auPRC score from ChromDL. Green indicates that ChromDL scored higher, and red indicates DanQ-JASPAR scored higher.

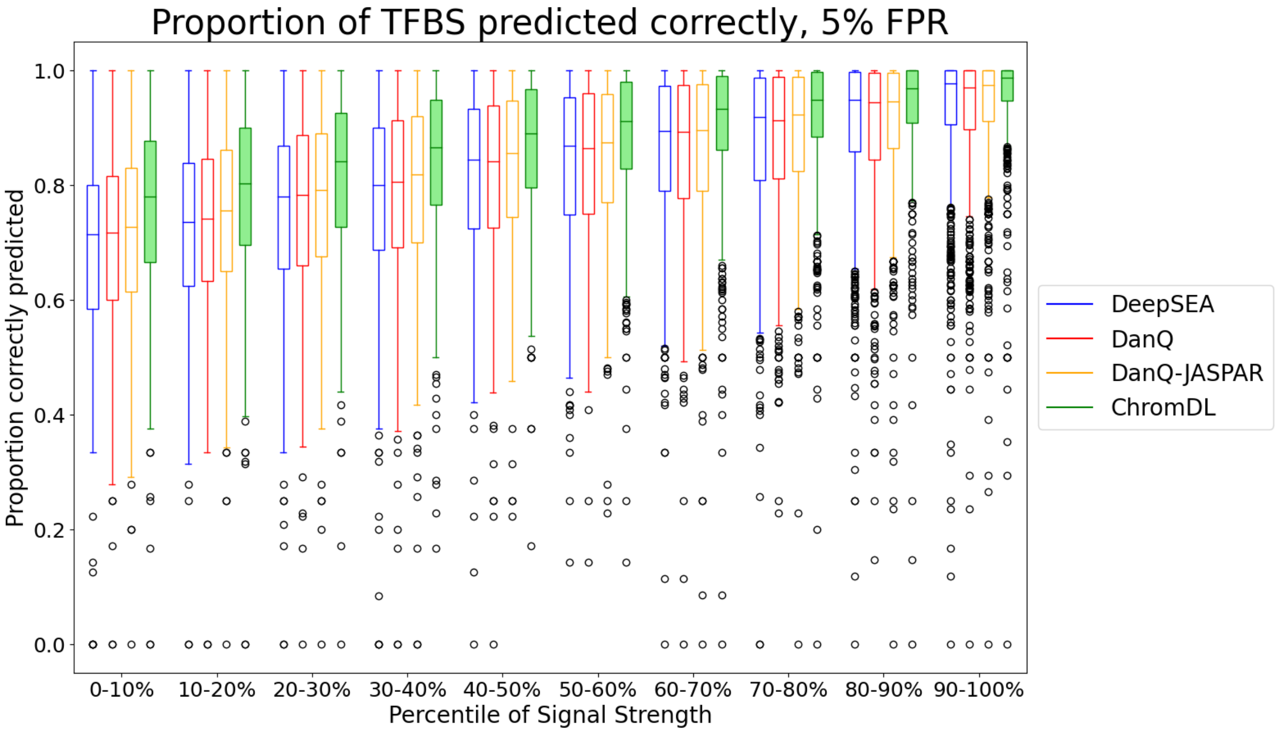


Figure 10: The proportion of TFBS ChIP-seq peaks in chromosomes 8 and 9 predicted correctly based on ChIP-seq signal strength, using 5% false positive rate thresholds calculated from the DeepSEA dataset.

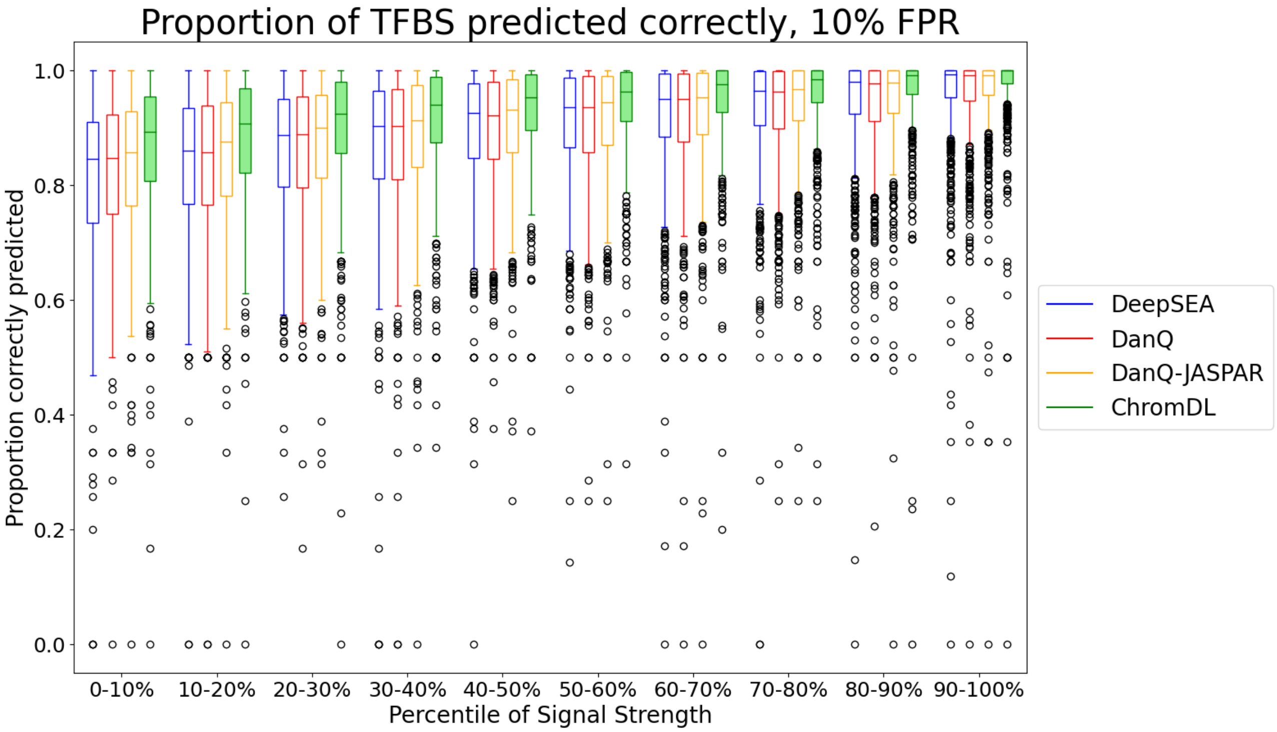


Figure 11: The proportion of TFBS ChIP-seq peaks in chromosomes 8 and 9 predicted correctly based on ChIP-seq signal strength, using 10% false positive rate thresholds calculated from the DeepSEA dataset.

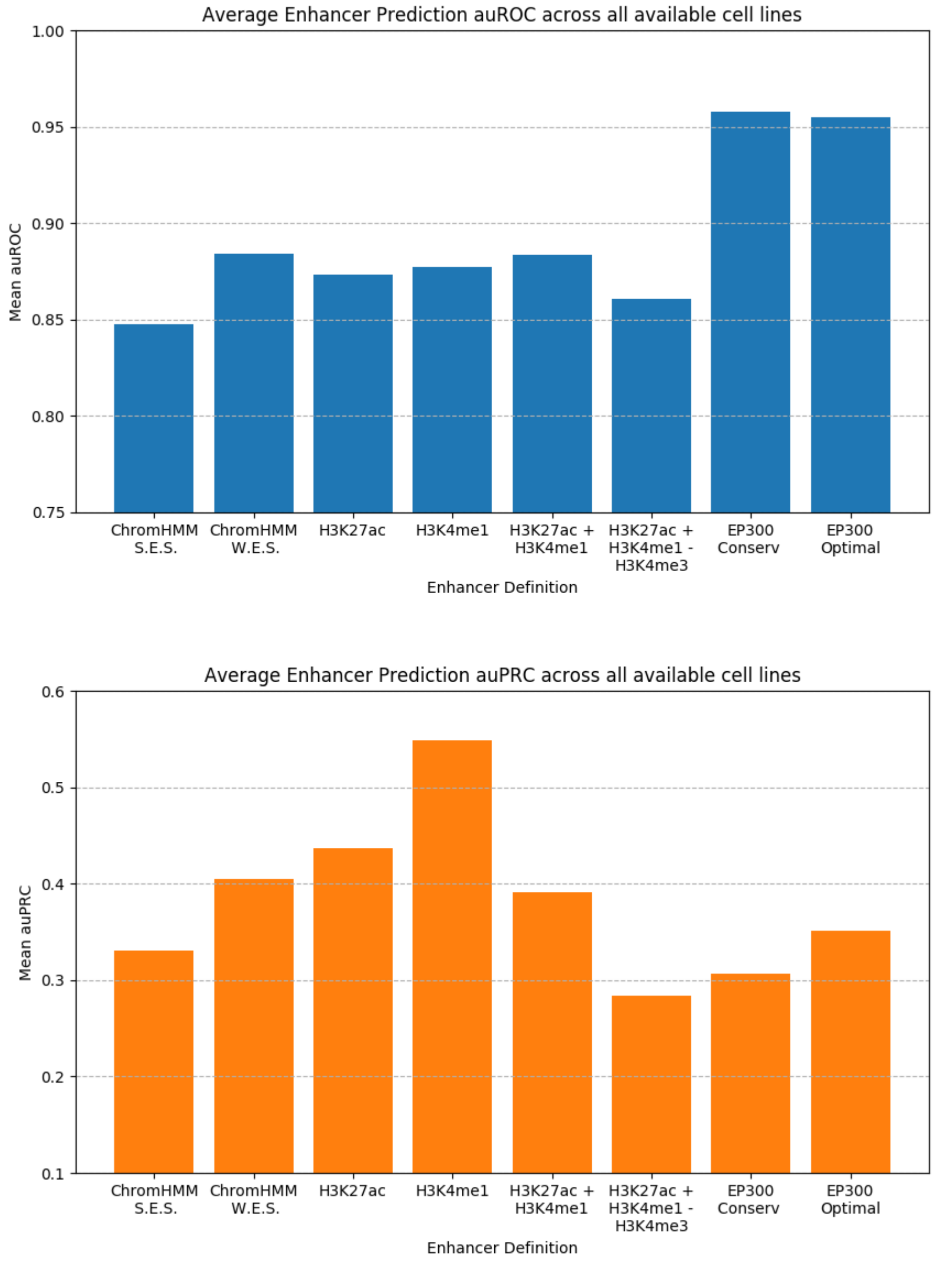


Figure 12: Average area under the receiver operating characteristic curve (auROC) and area under the precision recall curve (auPRC) metrics for the two-step ChromDL and TREDNet enhancer classifier in the prediction of all available cell line enhancers across the eight enhancer definitions.

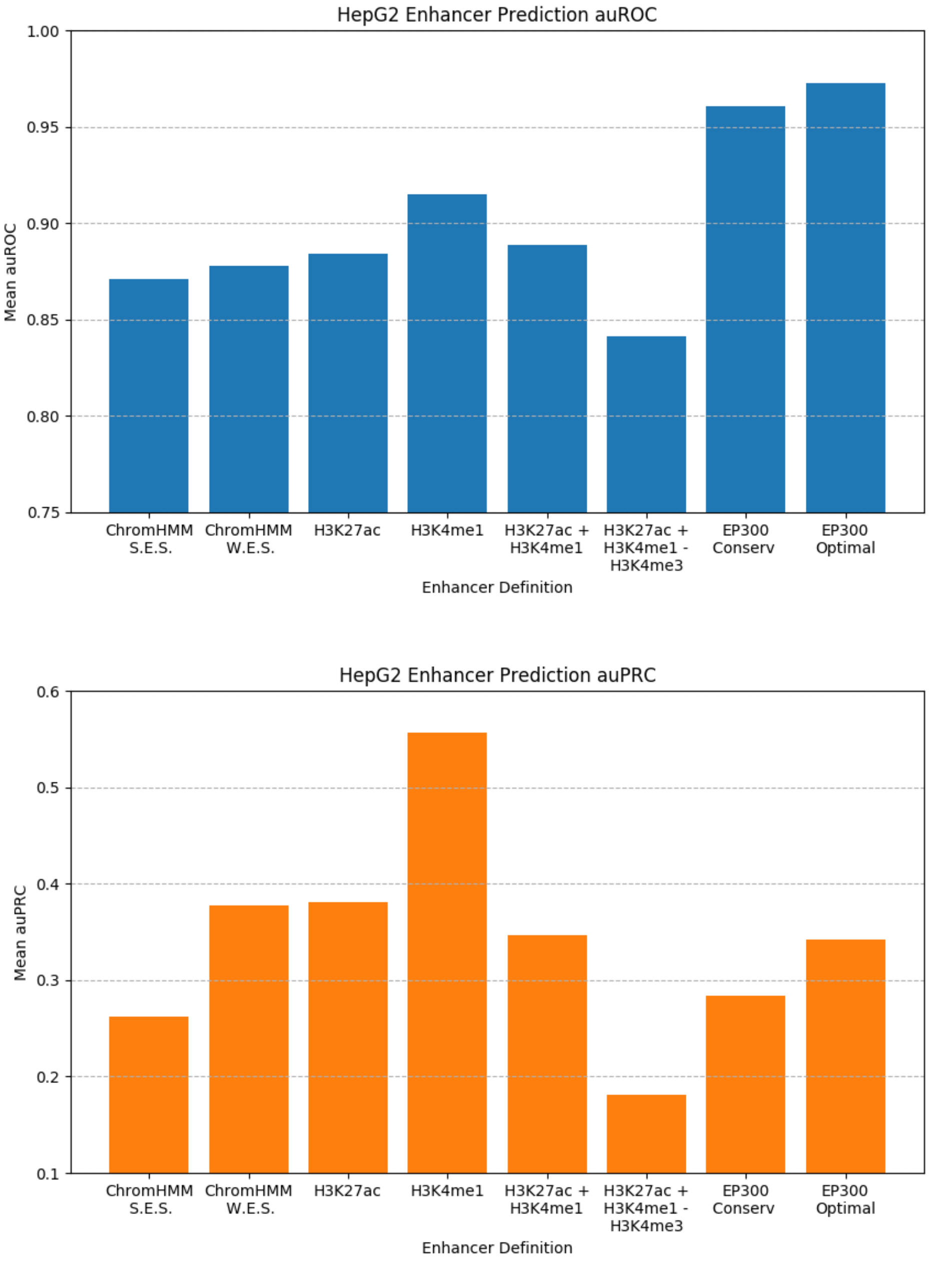


Figure 13: Area under the receiver operating characteristic curve (auROC) and area under the precision recall curve (auPRC) metrics for the two-step ChromDL and TREDNet enhancer classifier in the prediction of HepG2 enhancers across the eight enhancer definitions.

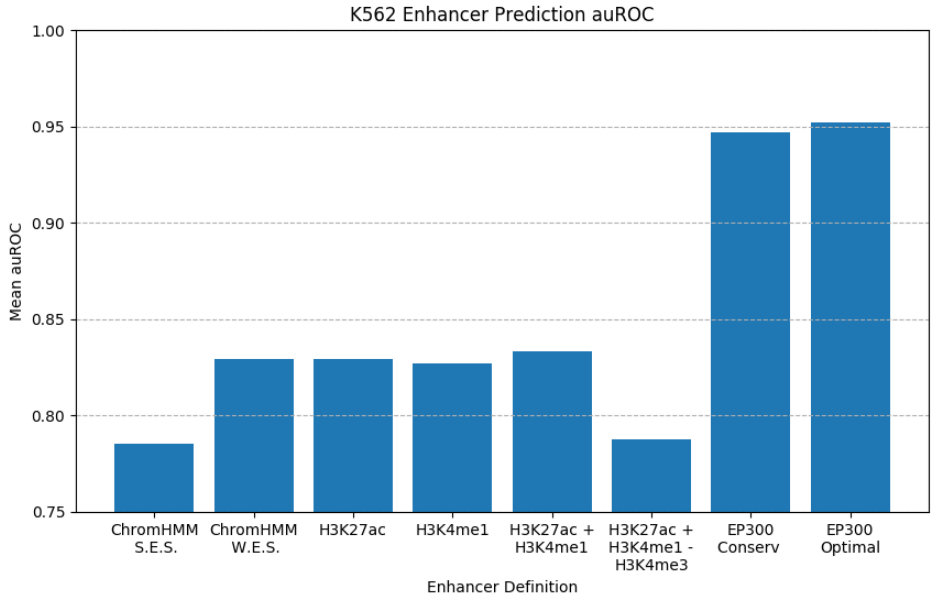


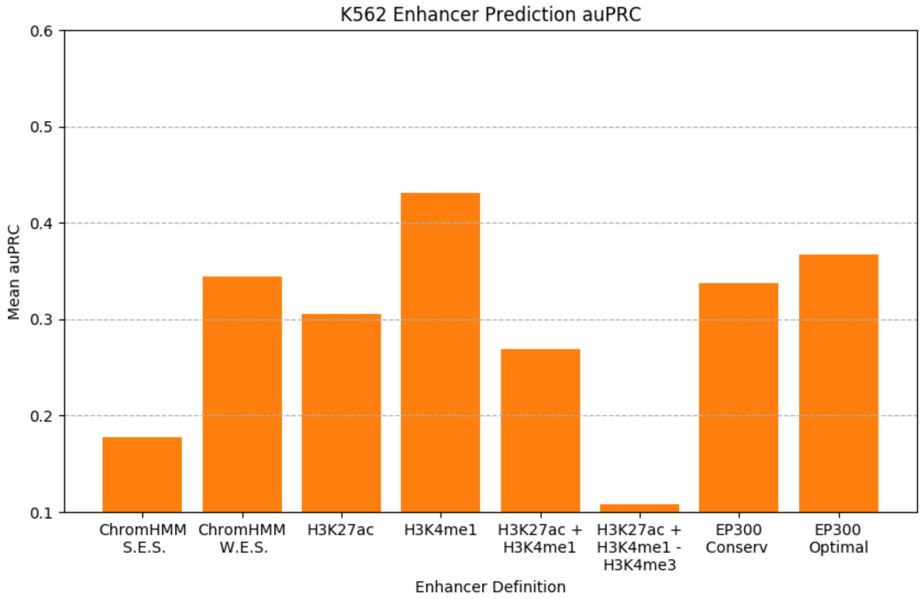


Figure 14: Area under the receiver operating characteristic curve (auROC) and area under the precision recall curve (auPRC) metrics for the two-step ChromDL and TREDNet enhancer classifier in the prediction of K562 enhancers across the eight enhancer definitions.

| Optimizer | auROC | auPRC |
|---|---|---|
| Adadelta | 0.6871246899 | 0.05599838184 |
| Adagrad | 0.7186713231 | 0.08439251365 |
| Adam* | 0.9475848281 | 0.4013451194 |
| Adamax | 0.9419850612 | 0.3836860648 |
| Ftrl | 0.5000027329 | 0.02057579854 |
| Nadam | N/A | N/A |
| RMSprop | N/A | N/A |
| SGD | 0.7475960588 | 0.09413193609 |

Table 1: Area under the receiver operating characteristic curve (auROC) and area under the precision recall curve (auPRC) metrics for ChromDL in trials run for 50 epochs using available Tensorflow optimizer algorithms. N/A signifies that the trial did not compile due to a memory error.

| | DeepSEA | DanQ | DanQ-JASPAR |
|---|---|---|---|
| All (919) | 883 (0.9608) | 745 (0.8107) | 707 (0.7693) |
| TFBS (690) | 677 (0.9812) | 582 (0.8435) | 551 (0.7986) |
| DHS (125) | 125 (1.0) | 124 (0.992) | 122 (0.976) |
| HM (104) | 81 (0.7788) | 39 (0.375) | 34 (0.3269) |

Table 2: The number and proportion of labels for each of the four categories (All, TFBS, DHS, and HM) that ChromDL has a higher area under the precision recall curve (auPRC) in a label by label comparison with each model.

| **Layer Removed** | **auROC** | **auPRC** |
|---|---|---|
| None | 0.9475848281 | 0.4013451194 |
| 1 - Bidirectional GRU | 0.9374964867 | 0.3640885309 |
| 2 - Separable Convolution | 0.944531268 | 0.3867219832 |
| 3 - Standard Convolution | 0.9454258992 | 0.3921503453 |
| 4 - Max Pooling | N/A | N/A |
| 5 - Bidirectional LSTM | 0.9462380911 | 0.3941711105 |
| 6 - Dropout (0.2) | 0.9446306885 | 0.3889063717 |
| 7 - Batch Normalization | 0.9459035395 | 0.3968426854 |
| 8 - Average Pooling | N/A | N/A |
| 9 - Bidirectional LSTM | 0.9446003652 | 0.3910669045 |

Table 3: Area under the receiver operating characteristic curve (auROC) and area under the precision recall curve (auPRC) metrics for ChromDL in trials where one of the nine removable layers was omitted. N/A signifies that the trial did not compile due to a memory error.

| | DeepSEA | | DanQ | | DanQ-JASPAR | |
|---|---|---|---|---|---|---|
| Bin | T-test | P-value | T-test | P-value | T-test | P-value |
| 0-10% | -8.1530773 | 7.90E-16 | -6.3276135 | 3.36E-10 | -4.9962495 | 6.59E-07 |
| 10-20% | -7.6469412 | 3.84E-14 | -6.7612259 | 2.02E-11 | -5.4985192 | 4.56E-08 |
| 20-30% | -7.969387 | 3.32E-15 | -6.8283979 | 1.28E-11 | -5.5361538 | 3.70E-08 |
| 30-40% | -8.0777258 | 1.43E-15 | -6.744457 | 2.25E-11 | -5.7229041 | 1.28E-08 |
| 40-50% | -7.1021338 | 1.97E-12 | -6.5205906 | 9.81E-11 | -5.5219765 | 4.00E-08 |
| 50-60% | -7.5988831 | 5.51E-14 | -6.7026998 | 2.98E-11 | -5.6250006 | 2.25E-08 |
| 60-70% | -6.8399662 | 1.19E-11 | -6.9661814 | 5.04E-12 | -5.8300435 | 6.90E-09 |
| 70-80% | -6.6332441 | 4.71E-11 | -6.557047 | 7.76E-11 | -5.4820492 | 5.00E-08 |
| 80-90% | -6.3759487 | 2.48E-10 | -6.6762581 | 3.55E-11 | -5.5889469 | 2.75E-08 |
| 90-100% | -6.1561746 | 9.77E-10 | -6.7621986 | 2.01E-11 | -5.3216468 | 1.20E-07 |

Table 4: One sided T-test statistics for ChromDL having a higher average proportion of TFBS peaks predicted correctly than the comparison models, 3% FPR.

| | DeepSEA | | DanQ | | DanQ-JASPAR | |
|---|---|---|---|---|---|---|
| Bin | T-test | P-value | T-test | P-value | T-test | P-value |
| 0-10% | -8.1652124 | 7.18E-16 | -6.8428256 | 1.16E-11 | -5.2912172 | 1.41E-07 |
| 10-20% | -7.8132325 | 1.10E-14 | -7.1254394 | 1.67E-12 | -5.4548504 | 5.80E-08 |
| 20-30% | -8.0607621 | 1.63E-15 | -6.7690846 | 1.91E-11 | -5.627678 | 2.21E-08 |
| 30-40% | -7.8062037 | 1.16E-14 | -7.0208603 | 3.45E-12 | -5.7098788 | 1.38E-08 |
| 40-50% | -7.4687997 | 1.43E-13 | -7.3002314 | 4.84E-13 | -5.8273272 | 7.01E-09 |
| 50-60% | -7.5769655 | 6.48E-14 | -6.9592396 | 5.29E-12 | -5.6120398 | 2.42E-08 |
| 60-70% | -6.6709247 | 3.68E-11 | -6.8308707 | 1.27E-11 | -5.6788849 | 1.65E-08 |
| 70-80% | -6.450614 | 1.54E-10 | -6.557902 | 7.71E-11 | -5.275569 | 1.54E-07 |
| 80-90% | -5.635925 | 2.11E-08 | -6.3053207 | 3.87E-10 | -4.9235979 | 9.53E-07 |
| 90-100% | -5.5641251 | 3.16E-08 | -6.3029066 | 3.93E-10 | -4.4592625 | 8.90E-06 |

Table 5: One sided T-test statistics for ChromDL having a higher average proportion of TFBS peaks predicted correctly than the comparison models, 5% FPR.

| | DeepSEA | | DanQ | | DanQ-JASPAR | |
|---|---|---|---|---|---|---|
| Bin | T-test | P-value | T-test | P-value | T-test | P-value |
| 0-10% | -7.3376966 | 3.70E-13 | -6.3272608 | 3.37E-10 | -4.822893 | 1.57E-06 |
| 10-20% | -7.1495413 | 1.41E-12 | -6.6367362 | 4.60E-11 | -5.3132669 | 1.25E-07 |
| 20-30% | -7.1874535 | 1.08E-12 | -6.9036245 | 7.71E-12 | -5.0893623 | 4.09E-07 |
| 30-40% | -7.0427247 | 2.97E-12 | -6.8057454 | 1.50E-11 | -4.9875194 | 6.89E-07 |
| 40-50% | -6.975797 | 4.71E-12 | -7.2603887 | 6.44E-13 | -5.3713821 | 9.17E-08 |
| 50-60% | -6.9271773 | 6.58E-12 | -7.086537 | 2.19E-12 | -5.3643086 | 9.53E-08 |
| 60-70% | -6.3918741 | 2.24E-10 | -6.330846 | 3.30E-10 | -5.4060833 | 7.59E-08 |
| 70-80% | -5.4230928 | 6.92E-08 | -5.9986442 | 2.54E-09 | -4.5047111 | 7.21E-06 |
| 80-90% | -4.7781734 | 1.96E-06 | -5.5370904 | 3.68E-08 | -4.1382798 | 3.71E-05 |
| 90-100% | -4.1616656 | 3.36E-05 | -4.6429692 | 3.76E-06 | -3.1909802 | 1.45E-03 |

Table 6: One sided T-test statistics for ChromDL having a higher average proportion of TFBS peaks predicted correctly than the comparison models, 10% FPR.

| Cell line | Query ID | Optimal Offset | p-value | E-value | q-value | Overlap | Query consensus | Target consensus | Ori. |
|---|---|---|---|---|---|---|---|---|---|
| | DeepSEA | 2 | 4.68E-08 | 4.68E-08 | 9.35E-08 | 10 | ATGACTCATC | GGATGACTCATTCC | - |
| H1-hESC | ChromDL | 3 | 1.07E-07 | 1.07E-07 | 2.13E-07 | 10 | GTGAGTCATC | GGAATGAGTCATCC | + |
| c-Jun | DanQ-JASPAR | 0 | 1.50E-07 | 1.50E-07 | 3.00E-07 | 12 | GGGTGAGTCATC | GGATGACTCATTCC | - |
| | DanQ | 3 | 2.35E-07 | 2.35E-07 | 2.35E-07 | 10 | ATGACTCACC | GGAATGAGTCATCC | + |
| | DanQ-JASPAR | 1 | 5.94E-10 | 5.94E-10 | 1.19E-09 | 12 | GGATGAGTCATC | GGAATGAGTCATCC | + |
| HeLa-S3 | ChromDL | 1 | 7.16E-10 | 7.16E-10 | 1.43E-09 | 12 | CGATGAGTCATC | GGAATGAGTCATCC | + |
| c-Fos | DanQ | 1 | 7.64E-10 | 7.64E-10 | 1.53E-09 | 12 | GGATGAGTCATC | GGAATGAGTCATCC | + |
| | DeepSEA | 1 | 9.11E-10 | 9.11E-10 | 1.82E-09 | 12 | GGATGAGTCATC | GGAATGAGTCATCC | + |
| | DanQ-JASPAR | 1 | 2.07E-09 | 2.07E-09 | 3.32E-09 | 12 | GGATGAGTCATC | GGAATGAGTCATCC | + |
| HeLa-S3 | ChromDL | 0 | 5.58E-09 | 5.58E-09 | 1.12E-08 | 12 | GGATGAGTCATC | GGATGACTCATTCC | - |
| c-Jun | DanQ | 0 | 1.64E-08 | 1.64E-08 | 3.29E-08 | 12 | CGGTGACTCACC | GGATGACTCATTCC | - |
| | DeepSEA | 3 | 3.78E-08 | 3.78E-08 | 6.97E-08 | 10 | ATGAGTCATC | GGAATGAGTCATCC | + |
| | ChromDL | 1 | 1.01E-09 | 1.01E-09 | 2.01E-09 | 12 | TGATGAGTCATC | GGAATGAGTCATCC | + |
| HepG2 | DanQ-JASPAR | 3 | 4.13E-08 | 4.13E-08 | 4.79E-08 | 10 | ATGACTCATC | GGAATGAGTCATCC | + |
| c-Jun | DanQ | 3 | 5.32E-08 | 5.32E-08 | 8.16E-08 | 10 | ATGACTCATC | GGAATGAGTCATCC | + |
| | DeepSEA | 3 | 8.09E-08 | 8.09E-08 | 9.29E-08 | 10 | ATGACTCATC | GGAATGAGTCATCC | + |
| | ChromDL | 0 | 1.55E-09 | 1.55E-09 | 3.10E-09 | 12 | AGATGACTCATT | GGATGACTCATTCC | - |
| HUVEC | DanQ-JASPAR | 0 | 2.57E-09 | 2.57E-09 | 5.15E-09 | 12 | AGATGACTCATT | GGATGACTCATTCC | - |
| c-Fos | DanQ | 0 | 3.05E-09 | 3.05E-09 | 6.10E-09 | 12 | GGATGAGTCATC | GGATGACTCATTCC | - |
| | DeepSEA | 0 | 7.62E-09 | 7.62E-09 | 8.83E-09 | 12 | GGATGAGTCATC | GGATGACTCATTCC | - |
| | ChromDL | 0 | 1.04E-09 | 1.04E-09 | 2.08E-09 | 12 | GGATGACTCATT | GGATGACTCATTCC | - |
| HUVEC | DanQ | 0 | 3.75E-09 | 3.75E-09 | 7.49E-09 | 12 | AAATGACTCATT | GGATGACTCATTCC | - |
| c-Jun | DeepSEA | 3 | 1.89E-05 | 1.89E-05 | 3.77E-05 | 10 | ATGAGTCATC | GGAATGAGTCATCC | + |
| | DanQ-JASPAR | 3 | 2.49E-05 | 2.49E-05 | 2.49E-05 | 10 | ATGAGTCATC | GGAATGAGTCATCC | + |
| | DanQ | 1 | 3.53E-09 | 3.53E-09 | 7.07E-09 | 12 | GGATGAGTCATC | GGAATGAGTCATCC | + |
| K562 | DeepSEA | 1 | 5.02E-08 | 5.02E-08 | 1.00E-07 | 12 | CGATGAGTCATC | GGAATGAGTCATCC | + |
| c-Fos | ChromDL | 1 | 1.01E-07 | 1.01E-07 | 2.01E-07 | 12 | GGTGAGTCATCC | GGATGACTCATTCC | - |
| | DanQ-JASPAR | 3 | 4.42E-07 | 4.42E-07 | 7.57E-07 | 8 | ATGAGTCA | GGAATGAGTCATCC | + |
| | ChromDL | 1 | 3.02E-08 | 3.02E-08 | 6.04E-08 | 12 | GGATGAGTCATC | GGAATGAGTCATCC | + |
| K562 | DeepSEA | 3 | 4.81E-08 | 4.81E-08 | 9.61E-08 | 10 | ATGAGTCATC | GGAATGAGTCATCC | + |
| c-Jun | DanQ | 1 | 5.62E-08 | 5.62E-08 | 1.12E-07 | 10 | GATGAGTCAT | GGATGACTCATTCC | - |
| | DanQ-JASPAR | 3 | 4.33E-07 | 4.33E-07 | 8.66E-07 | 10 | ATGACTCATC | GGAATGAGTCATCC | + |

Table 7: TOMTOM metrics when mapping *de novo* generated motifs of ChromDL, DeepSEA, DanQ, and DanQ-JASPAR using HOMER to published Protein Binding Microarray position weight matrices.

| Cell line | Query ID | Optimal Offset | p-value | E-value | q-value | Overlap | Query consensus | Target consensus | Ori. |
|---|---|---|---|---|---|---|---|---|---|
| H1-hESC c-Jun | ChromDL | 1 | 1.47E-07 | 1.47E-07 | 2.95E-07 | 13 | GATGACTCATCCCTCTGGCA | GGATGACTCATTCC | - |
| | DanQ-JASPAR | 1 | 2.09E-07 | 2.09E-07 | 4.18E-07 | 13 | GATGACTCACCCCTTTGGCA | GGATGACTCATTCC | - |
| | DanQ | 1 | 8.19E-08 | 8.19E-08 | 1.64E-07 | 13 | GATGACTCACCCCTTTGGCAT | GGATGACTCATTCC | - |
| | DeepSEA | 1 | 8.34E-08 | 8.34E-08 | 1.67E-07 | 13 | GATGACTCACCCCTTTGGCAT | GGATGACTCATTCC | - |
| HeLa-S3 c-Fos | ChromDL | 0 | 2.07E-09 | 2.07E-09 | 4.14E-09 | 14 | CCAGTGACTCATCCT | GGAATGAGTCATCC | + |
| | DanQ-JASPAR | 3 | 6.45E-08 | 6.45E-08 | 8.50E-08 | 11 | ATGAGTCATCC | GGAATGAGTCATCC | + |
| | DanQ | 1 | 7.15E-08 | 7.15E-08 | 1.43E-07 | 11 | TATGAGTCACT | GGATGACTCATTCC | - |
| | DeepSEA | 1 | 7.66E-08 | 7.66E-08 | 1.53E-07 | 11 | TATGAGTCACT | GGATGACTCATTCC | - |
| HeLa-S3 c-Jun | DeepSEA | 0 | 3.53E-10 | 3.53E-10 | 7.07E-10 | 14 | GCAATGAGTCATCA | GGAATGAGTCATCC | + |
| | ChromDL | 2 | 7.05E-08 | 7.05E-08 | 1.41E-07 | 11 | AGTGACTCATC | GGAATGAGTCATCC | + |
| | DanQ-JASPAR | 1 | 1.19E-07 | 1.19E-07 | 1.36E-07 | 11 | GGATGACTCAT | GGAATGAGTCATCC | + |
| | DanQ | 1 | 1.98E-07 | 1.98E-07 | 3.23E-07 | 11 | GATGACTCATC | GGATGACTCATTCC | - |
| HepG2 c-Jun | ChromDL | 0 | 1.21E-10 | 1.21E-10 | 2.43E-10 | 14 | TGATGACTCATCTT | GGATGACTCATTCC | - |
| | DanQ-JASPAR | 1 | 1.61E-07 | 1.61E-07 | 3.22E-07 | 11 | GATGACTCATC | GGATGACTCATTCC | - |
| | DanQ | 1 | 1.93E-07 | 1.93E-07 | 3.00E-07 | 11 | GATGACTCATC | GGATGACTCATTCC | - |
| | DeepSEA | 1 | 2.02E-07 | 2.02E-07 | 3.47E-07 | 11 | GATGACTCATC | GGATGACTCATTCC | - |
| HUVEC c-Fos | DanQ | -1 | 4.19E-09 | 4.19E-09 | 8.37E-09 | 14 | GAAGATGAGTCATAG | GGAATGAGTCATCC | + |
| | DanQ-JASPAR | -2 | 4.29E-08 | 4.29E-08 | 8.57E-08 | 12 | AGGGAATGAGTCAT | GGAATGAGTCATCC | + |
| | DeepSEA | 2 | 5.71E-08 | 5.71E-08 | 1.14E-07 | 11 | ATGAGTCATTC | GGATGACTCATTCC | - |
| | ChromDL | 2 | 9.08E-08 | 9.08E-08 | 1.82E-07 | 11 | ATGAGTCACCC | GGATGACTCATTCC | - |
| HUVEC c-Jun | DanQ-JASPAR | 0 | 5.24E-10 | 5.24E-10 | 1.05E-09 | 14 | GTATGAGTCATTTCC | GGATGACTCATTCC | - |
| | DeepSEA | 2 | 4.74E-08 | 4.74E-08 | 8.48E-08 | 11 | ATGACTCATCC | GGATGACTCATTCC | - |
| | DanQ | 2 | 7.09E-08 | 7.09E-08 | 8.22E-08 | 11 | ATGACTCATCC | GGATGACTCATTCC | - |
| | ChromDL | 2 | 1.28E-07 | 1.28E-07 | 2.19E-07 | 11 | AATGAGTCATC | GGAATGAGTCATCC | + |
| K562 c-Fos | DanQ | 1 | 1.32E-07 | 1.32E-07 | 2.49E-07 | 11 | GGTGACTCATG | GGATGACTCATTCC | - |
| | ChromDL | 1 | 7.59E-07 | 7.59E-07 | 1.52E-06 | 10 | GATGACTCAT | GGATGACTCATTCC | - |
| | DanQ-JASPAR | 1 | 1.08E-06 | 1.08E-06 | 2.17E-06 | 10 | GATGACTCAT | GGATGACTCATTCC | - |
| | DeepSEA | 1 | 1.40E-06 | 1.40E-06 | 2.80E-06 | 10 | GATGACTCAG | GGATGACTCATTCC | - |
| K562 c-Jun | ChromDL | -1 | 3.88E-09 | 3.88E-09 | 7.75E-09 | 14 | AATGATGACTCATAG | GGAATGAGTCATCC | + |
| | DanQ | -1 | 4.75E-09 | 4.75E-09 | 9.51E-09 | 14 | AAGGATGACTCATAC | GGAATGAGTCATCC | + |
| | DanQ-JASPAR | 1 | 6.71E-07 | 6.71E-07 | 1.34E-06 | 11 | GATGACTCATC | GGATGACTCATTCC | - |
| | DeepSEA | 1 | 8.77E-07 | 8.77E-07 | 1.45E-06 | 11 | GATGACTCATC | GGATGACTCATTCC | - |

Table 8: TOMTOM metrics when mapping *de novo* generated motifs of ChromDL, DeepSEA, DanQ, and DanQ-JASPAR using MEME to published Protein Binding Microarray position weight matrices.
